# An ALS-associated TARDBP mutation drives cryptic exon inclusion and RNA dysregulation

**DOI:** 10.1101/2025.11.09.687455

**Authors:** Qi Zhang, Meng Liu, Xiaojuan Fan, Natalie Chin, Yue Xu, Jessica Suh, Soukaina Amniouel, James Virga, Brothely Jones, Kaari Linask, Jizhong Zou, Christopher Grunseich, Markus Hafner, Lichun Ma, Yihong Ye, Wei Zheng

**Affiliations:** Therapeutic Development Branch, National Center for Advancing Translational Sciences, National Institutes of Health, Bethesda, MD, 20892, USA; Cancer Data Science laboratory, National Cancer Institute, National Institutes of Health, Bethesda, MD, 20892, USA; RNA Molecular Biology Laboratory, National Institute of Arthritis and Musculoskeletal and Skin Disease, National Institutes of Health, Bethesda, MD, 20892, USA; Laboratory of Molecular Biology, National Institute of Diabetes, Digestive, and Kidney Diseases, National Institutes of Health, Bethesda, MD, 20892, USA; iPSC Core, National Heart, Lung, and Blood Institute, National Institutes of Health, Bethesda, MD, 20892, USA; Neurogenetics Branch, National Institute of Neurological Disorders and Stroke, National Institutes of Health, Bethesda, MD, 20892, USA; Liver Disease Branch, National Institute of Diabetes, Digestive, and Kidney Diseases, National Institutes of Health, Bethesda, MD, 20892, USA; Chinese Institute for Brain Research, Chinese Academy of Medical Sciences, Changping, Beijing 102206

**Keywords:** TDP-43 proteinopathies, RNA splicing, cryptic exon, iPSC-derived organoid, PRDM2, oxidative stress

## Abstract

TAR DNA–binding protein 43 (TDP-43) proteinopathy is a defining pathological feature of amyotrophic lateral sclerosis (ALS) and frontotemporal dementia (FTD), yet the downstream molecular events linking TDP-43 dysfunction to neurodegeneration remain incompletely understood. Here, we model TDP-43 proteinopathy by introducing the ALS/FTD-associated TARDBP K181E mutation into human induced pluripotent stem cells and differentiating them into three-dimensional forebrain organoids. Organoids carrying the K181E mutation exhibit spontaneous TDP-43 hyperphosphorylation, cytoplasmic p-TDP43 accumulation, RNA dysregulation, pro-inflammatory signaling, and activation of apoptotic pathways without TDP-43 overexpression or external stress. Single-cell transcriptomics and enhanced crosslinking immunoprecipitation reveal that the K181E mutation alters TDP-43 RNA-binding specificity, causing widespread RNA mis-splicing including cryptic exon inclusion. Among affected targets, we identify cryptic exon inclusion in the transcription factor *PRDM2*. PRDM2 cryptic exon-derived peptide immunoreactivity was detected in ALS spinal motor neurons and associated with p-TDP-43 pathology. These findings identify PRDM2 mis-splicing as a candidate downstream event linking TDP-43 dysfunction relevant to ALS and FTD.

**One sentence summary:** Human brain organoids reveal how mutant TDP-43 disrupts RNA processing and causes neuronal stress.

## Introduction

The neurodegenerative disorders amyotrophic lateral sclerosis (ALS) and frontotemporal dementia (FTD) are united by the pathological mis-localization and aggregation of TAR DNA-binding protein 43 (TDP-43) in affected neurons and glial cells [1, 2]. TDP-43 aggregation is present in approximately 97% of ALS and 45% of FTD cases, as well as in subsets of other neurodegenerative conditions including Alzheimer’s disease and chronic traumatic encephalopathy. These diseases are now collectively termed TDP-43 proteinopathies [1, 3]. Despite extensive research, the cellular and molecular mechanisms underpinning TDP-43 aggregation and its contribution to neurodegeneration remain poorly understood.

TDP-43 is a nuclear RNA-binding protein with essential roles in RNA metabolism, including RNA splicing, transport and stability [4, 5]. It contains an N-terminal dimerization domain, two RNA-recognition motifs (RRM1 and RRM2), and a C-terminal low-complexity domain (LCD) [6]. The LCD of TDP-43 is known to undergo liquid-liquid phase separation (LLPS) [6, 7]; Under pathological or stress conditions, this property is altered such that TDP-43 undergoes a liquid-to-solid phase transition [7, 8]. This leads to abnormal nuclear RNA regulation while generating cytoplasmic p-TDP-43 accumulation [4, 6, 9]. Notably, existing experimental models often rely on artificial manipulations such as TDP-43 overexpression or stress induction [10–14]. While these models have provided valuable insights into the potential disease mechanisms, they often fail to recapitulate endogenous p-TDP43 accumulation and other disease-associated pathologies, and are therefore, not ideal for therapeutic evaluation or mechanistic study.

Recent technology advancement in induced pluripotent stem cells (iPSCs) and CRISPR-mediated gene editing, combined with the establishment of three-dimensional (3D) organoid models have provided new opportunities for disease modeling [15–17]. These 3D cell assemblies, derived from CRISPR-engineered iPSCs bearing endogenous disease mutations, generate distinct cell types that self-assemble into brain-like structures with extensive cell-cell interactions. Thus, organoids better mimic the cellular complexity and organization of the human brain.

To date, ∼70 genetic mutations in *TARDBP* have been linked to ALS or FTD. While the majority of ALS/FTD-associated mutations are located within the LCD, many are located in or near the RRMs. These TDP-43 mutants are often defective in RNA-binding [6], which indirectly affect their biochemical properties in LLPS or liquid-solid regulation [18]. Of particular interest is a recently identified severe mutation (K181E) reported in a family with ALS/FTD, in which a 38-year-old man developed ALS and survived for 36 months after symptom onset, whereas his 76-year-old father developed symptoms of both ALS and FTD and survived for 80 months after symptom onset [19]. The TDP-43 K181E mutant cannot bind the canonical TDP-43 binding sequence in RNA, yet it can still undergo LLPS in vitro [20, 21]. In cells, overexpressed TDP-43 K181E mutant is hyperphosphorylated and forms large irregularly shaped aggregates without external stress [19]. However, it is unclear how this mutation causes neuronal cell dysfunction.

Since patient cells are unavailable, we introduced the TARDBP K181E mutation into a genetically defined neuronal platform to examine downstream consequences of impaired TDP-43 RNA processing under endogenous conditions [22, 23]. We differentiated these cells into neurons or 3-D brain-like organoids and found that 3-D organoids bearing the disease mutation display spontaneous TDP-43 phosphorylation, cytoplasmic accumulation and nuclear TDP-43 depletion. Our OMICs-based approach further revealed key molecular signatures in these organoids, closely resembling patient phenotypes including RNA dysregulation and neuroinflammation. Importantly, our study has linked mis-splicing of several mRNAs including the epigenetic PRDM2-encoding gene to TDP-43 dysfunction. We further validated that PRDM2 mis-splicing identified in the K181E organoid model generates an abnormal polypeptide in ALS patient spinal cord and tested the potential impact of reduced PRDM2 expression on neuronal homeostasis.

## Results

### The TARDBP K181E mutation induces spontaneous TDP-43 proteinopathy under endogenous expression conditions

To determine how the ALS-associated TARDBP K181E mutation affects TDP-43 function under endogenous expression conditions, we used CRISPR-mediated gene editing to introduce the K181E mutation at the endogenous *TARDBP* locus in a well-characterized dementia genetic risk-free iPSC line followed by extensive whole exome sequencing to rule out CRISPR mediated off target effect (see method and table S6). Genomic sequencing identified several K181E heterozygous and homozygous (HM) knock-in (KI) clones with normal karyotypes and confirmed pluripotency (Supplementary Fig. S1A, B). We transduced clones with a lentiviral vector that enabled their differentiation into excitatory neurons (Exci-neurons) via tetracycline-induced expression of neurogenin-2 (NGN2) [24]. We then used immunoblotting to examine hyperphosphorylated TDP-43 (p-TDP-43) at Ser409/410, a hallmark of TDP-43 proteinopathies by a widely used antibody [25–28], but failed to detect it in differentiated neurons under normal growth conditions regardless of the genotype (Supplementary Fig. S1C, D); Only after treatment with a proteasome inhibitor (MG132), did we observe a small amount of p-TDP-43 in both wild-type (WT) and K181E neurons, suggesting that p-TDP-43 generated from WT and K181E TDP-43 is rapidly degraded by the proteasome and that this 2D cell model does not phenocopy TDP-43 proteinopathies.

To generate a better TDP-43 proteinopathy model, we used a growth factor-based system to differentiate and grew these iPSC cells from multiple clones into 3D organoids (Fig. 1A) because organoids can reconstruct cell diversity and cell-cell communications essential for pathogenesis. To this end, we first screened and identified a ROCK2 inhibitor as a critical element that supported forebrain-like growth during the early phase of differentiation, particularly for cells bearing the disease mutation (Supplementary Fig. S1E-G). We then maintained the cultures for 85 days, at which point, we used immunostaining to confirm the robust expression of key markers including SATB2 (Exci-neuron) and GFAP (glial marker) (Fig. 1B). The staining also revealed the formation of distinct forebrain layers, mirroring the laminar organization of the brain (Fig. 1C). Consistent with the developmental nature of forebrain organoid, these cultures represent an early-stage human neural system and were therefore used to investigate proximal molecular consequences of TDP-43 dysfunction under endogenous expression conditions.

**Figure 1.**
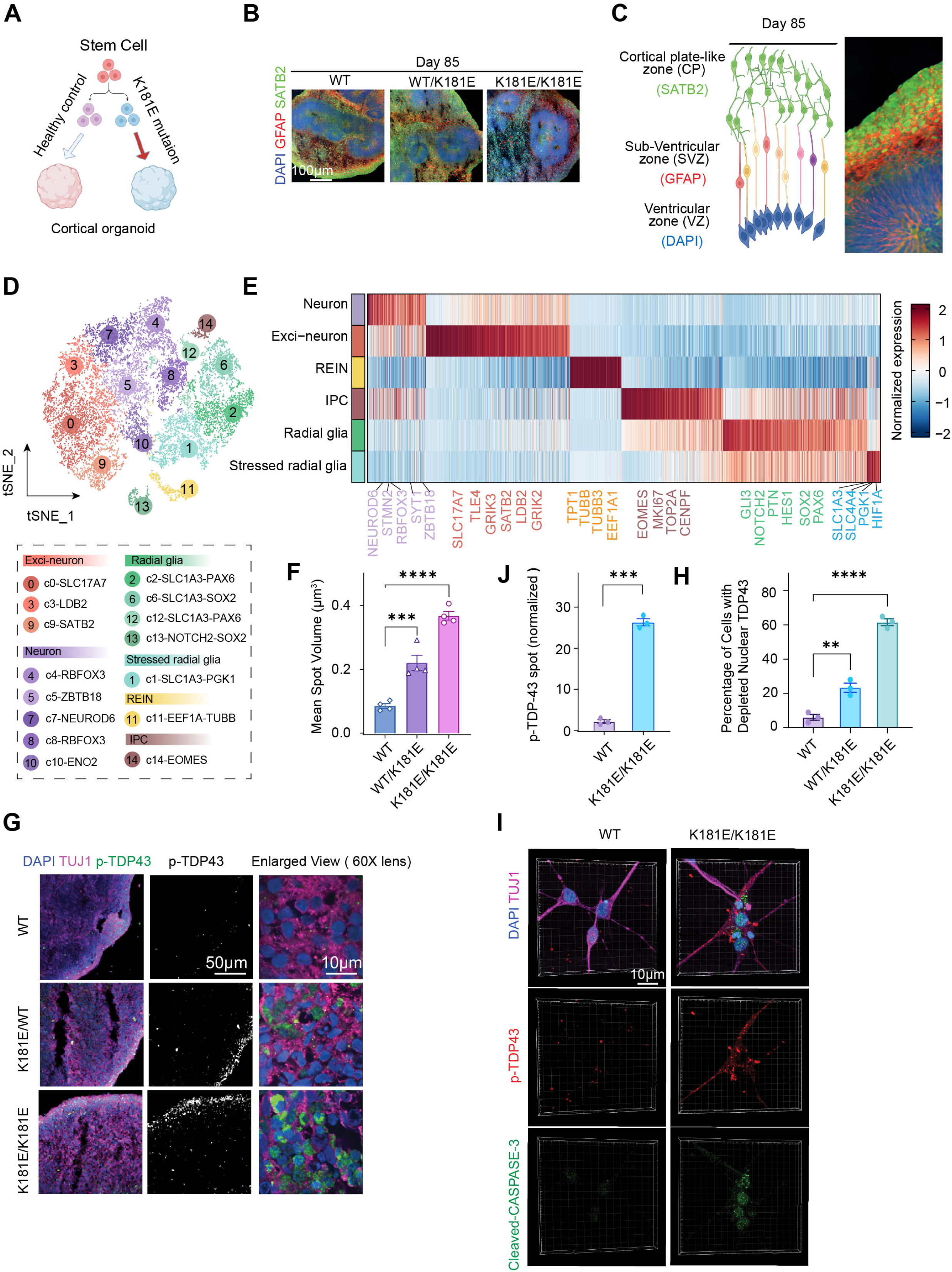
A forebrain brain organoid model of TDP-43 proteinopathies. **(A)** Schematic of forebrain brain organoids differentiation. **(B)** Immunostaining of organoid sections with antibodies against the indicated neuronal markers at day 85 ( n= 3-5 organoids, 2-3 iPSC clones, two individual batches). **(C)** Schematic and representative image showing a forebrain-like organization at day 85. **(D)** t-SNE visualization of 16549 single cells from WT and K181E/K181E organoids, colored by cluster. **(E)** Heatmap of cell type-specific genes (log2 fold change ≥0.5, adjusted P<0.05, expressed in ≥10% of cells); representative markers are grouped by cell type. **(F, G, H)** Phosphorylated TDP-43 accumulates in K181E mutant organoids. **(F)** Quantification of phosphorylated TDP-43 (p-TDP43) mean spot volume in 3D fluorescence images of immuno-stained organoids with the indicated genotypes. Error bars indicate mean ± s.e.m., **** p < 0.0001, *** p < 0.001 by one-way ANOVA, n=4 organoids. **(G)** Images were acquired from the comparable peripheral regions of organoid sections, which are enriched for cortical neuron-like cells under identical conditions. Enlarged views show the distribution of p-TDP-43 signal relative to DAPI-labeled nuclei and TUJ1-positive neuronal structures. Scale bar, 10 µm, n= 3-5 organoids from 2-3 iPSC clones, two individual batches. (**H**) Quantification of cells with depleted nuclear TDP43 with the indicated genotypes. Error bars indicate mean ± s.e.m., **** p < 0.0001, ** p < 0.01 by one-way ANOVA, n=3 organoids. **(I)** Representative images of neurons dissociated from mature organoids at day 107, cultured as isolated cells, and fixed at day 127. TUJ1, p-TDP-43, cleaved caspase-3, and DAPI are shown (n=3–5 organoids from 2–3 iPSC clones; two independent batches). (**J**) Quantification of normalized p-TDP-43 spot number per cell, calculated as detected p-TDP-43 spots divided by DAPI-positive nuclei per field. Data are mean ± SEM; unpaired Student’s t test; ***P<0.001; n=3 organoids.

Single-cell transcriptomic profiling of pooled organoids from two independent batches revealed various brain cell types including radial glia, stressed radial glia, intermediate progenitor cells (IPC), and Exci-neurons (Fig. 1D, E, Supplementary Fig. S2A-E, Supplementary Table S1), as reported previously [29] , indicating the formation of complex cellular structures in these organoids. The presence of both neuronal and radial glia underscores the physiological relevance of our model because abnormal neuron-glia interactions are known to induce neurodegeneration [16].

We next sectioned the organoids and performed immunostaining with antibodies against phosphorylated TDP-43 (p-TDP-43), focusing our analysis on the outer layers of the organoids that are enriched for cortical neurons. Noticeably, while minimal p-TDP-43 signal was detected in WT organoids, we saw some p-TDP-43 signal in heterozygous organoids and significantly more p-TDP-43 in K181E homozygous (HM) organoids suggesting an allele dose-dependent effect (see discussion). Immunostaining with TDP-43 antibodies showed that K181E HM organoids had a significant percentage of cells depleted of nuclear TDP43 accompanied by cytoplasmic p-TDP-43 accumulation (Fig. 1F-H, Supplementary Fig. S1H-I). Co-immunostaining using antibodies against the neuron specific protein TUJ1 showed substantial co-localization of p-TDP-43 with TUJ1, suggesting its accumulation in neurons in HM K181E organoids (Fig. 1G, right panels). However, our data does not exclude the possibility that other cell types may also exhibit TDP-43 pathology. Because HM K181E organoids had stronger TDP-43 proteinopathy phenotypes, we focused our study on HM K181E organoids.

p-TDP-43 signal in brain organoids displays large patch-like accumulation similar to that found in cortex tissue [30], obscuring its precise subcellular localization. To further confirm the cytosolic localization of p-TDP-43, we dissociated cells from the organoids and cultured them in monolayer. This allowed cell visualization with high resolution. As reported previously [12], most p-TDP-43 signals were detected in the cytoplasm in HM K181E cells (Fig. 1I, J). Moreover, staining with antibodies recognizing cleaved Caspase-3 detected Caspase-3 activation in some p-TDP-43 positive cells (Fig. 1I), consistent with the reported association of neurotoxicity with TDP-43 proteinopathy [31]. Thus, the 3D TDP-43 K181E organoid replicates key features of TDP-43 proteinopathy including endogenous TDP-43 cytoplasmic accumulation, nuclear TDP-43 depletion, and activation of cell apoptosis associated signaling in a subset of neurons. Importantly, these phenotypes were developed without TDP-43 overexpression or exposing cells to an external stressor.

### Transcriptomic characterization of TDP-43 K181E organoids

To evaluate the clinical relevance of our model, we first compared the overall dysregulated gene expression in HM K181E organoids with genes dysregulated in postmortem brain samples of dementia patients with confirmed TDP-43 proteinopathy [32]. Strikingly, among the 178 genes upregulated in patient samples [32], 177 (99.4%) were also upregulated in HM K181E organoids when compared to WT organoids. By contrast, among the 153 genes downregulated in patients, only 9 (5.9%) were similarly reduced in HM K181E organoids (Fig. 2A, Supplementary Table S2). Gene Ontology (GO) analysis of the 177 upregulated genes (Q1 in Fig. 2A) highlighted a significant enrichment in stress adaptation pathways such as protein translation, ribosomal biogenesis, and protein quality control (Fig. 2B). By contrast, genes down-regulated specifically in patients are mostly involved in synaptic connectivity regulations (e.g. glutamatergic synapse and gap junction) (Supplementary Fig. S3A). These cellular functions could be influenced by aging, inter-tissue communications, and other factors missing in organoids. We therefore conclude that our 3D organoid model recapitulates a subset of stress-associated transcriptional responses observed in TDP-43 proteinopathy patients.

**Figure 2.**
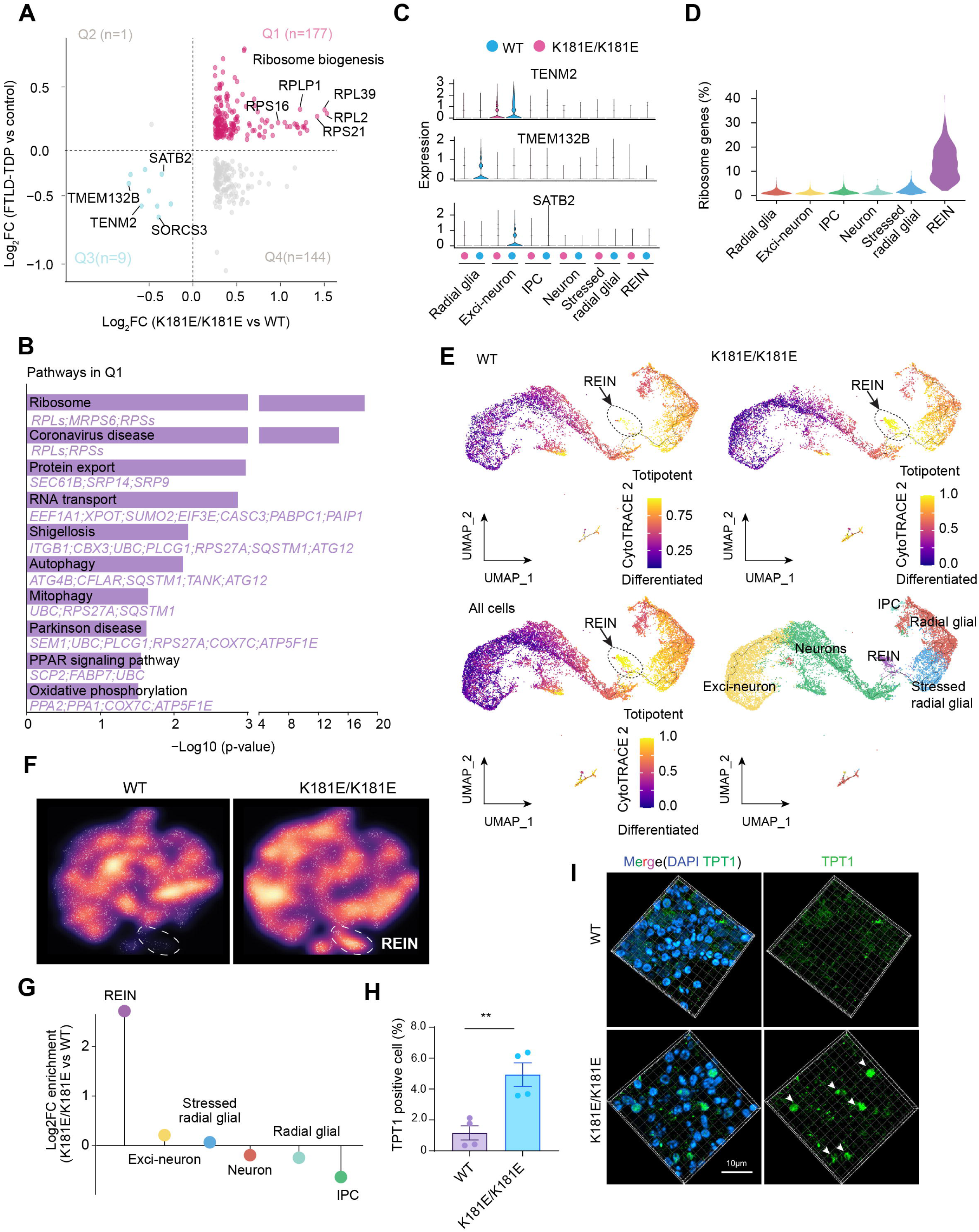
Single cell transcriptomic characterization of homozygous TDP-43 K181E organoids. **(A)** Comparison of differentially expressed genes in K181E/K181E versus WT forebrain organoids (x axis) and FTLD-TDP patient forebrain tissue versus control tissue (y axis), identifying 177 commonly up-regulated genes (Q1, magenta) and 9 commonly down-regulated genes (Q3, blue). **(B)** Top enriched KEGG pathways of the commonly upregulated genes in **(A)** (n=177) highlight the upregulation of protein biogenesis and quality control in mutant organoids. **(C)** Violin plots comparing the expression levels of TENM2, SATB2, and TMEM132B for the indicated cell types in WT and K181E/K181E forebrain organoids. **(D)** Violin plot showing the percentage of ribosome-related genes enriched in the indicated cell types. **(E)** CytoTRACE trajectory analysis of WT, K181E/K181E, and combined organoid cells. Higher scores indicate a less differentiated state. Ribo-enriched immature neurons (REIN), defined by coordinated up-regulation of ribosomal genes and neuronal identity markers, are outlined by dashed lines. Bottom right: UMAP of WT and K181E/K181E cells colored by cell type; black lines indicate graph structure. **(F)** Cell density plots showing the accumulation of Ribo-enriched immature neurons (REIN) (indicated by dashed lines) in K181E/K181E forebrain organoids. **(G)** Log_2_ FC of cell type enrichment in K181E/K181E forebrain organoids compared to WT organoids. **(H)** Quantification of TPT1 expression determined by immunostaining in **(I)** in organoids of the indicated genotypes. Error bars indicate mean ± s.e.m., ** p < 0.01 unpaired Student’s t-test, n=4 organoids. **(I)** Reconstructed 3D confocal fluorescence images of immuno-stained organoid sections show increased expression of TPT1 (green) in K181E/K181E organoids. Organoids were co-stained with DAPI (blue) to label nuclei. Scale bar, 10 µm, n= 3-5 organoids, 2-3 iPSC clones, two individual batches.

Cell type specific gene expression analysis showed that the commonly downregulated genes such as SATB2 and TENM2 had the most significantly reduced expression in Exci-neurons, while TMEM132B was more dramatically downregulated in radial glia in HM K181E mutant organoids (Fig. 2C). These genes have key functions in forebrain neuron development and synaptic plasticity [33–36]. Thus, their suppression may contribute to the abnormal brain functions in patients with TDP-43 proteinopathies.

Notably, scRNA-seq analysis revealed a distinct cluster of cells (∼5%) enriched in HM K181E organoids and characterized by coordinated upregulation of ribosomal and translation-related genes (Fig. 2D–G, Supplementary Table S1). This population expressed neuronal markers including SYT1, supporting a neuronal identity. Hierarchical analysis of transcriptomic profile across all cell types further indicated similar transcriptome of REIN population and excitatory neurons (Supplementary Fig. S3B). CytoTRACE2-based differentiation potency analysis suggested that these cells occupy a less differentiated neuronal state relative to other excitatory neuron populations (Fig. 2E). Based on this combined transcriptional profile, we refer to this population as ribo-enriched immature neurons (REINs).

Immunostaining of organoid sections using antibodies recognizing TPT1, a nuclear protein upregulated in REINs (Fig. 1E), confirmed the enrichment of this novel cell type in HM TDP-43 mutant organoids (Fig. 2H, I). Additional bioinformatic analyses of published patient RNA-seq datasets revealed enrichment of REIN-associated gene signatures in subsets of neurons from idiopathic ALS and FTD patients (Supplementary Fig. S3C), suggesting partial overlap with disease-relevant transcriptional signature [37]. However, because REIN-associated gene signature is also modestly upregulated in patient astrocytes and oligodendrocytes, these gene expression signatures alone do not identify REINs in ALS/FTD (see discussion). Although our study did not measure directly changes in translational activity, our analysis suggests highly expressed translational related genes corroborate recent findings that implicate abnormal protein translation in TDP-43 proteinopathies and related diseases [38].

### Single cell analysis reveals neuroinflammatory signatures in TDP-43 K181E organoids

To elucidate if TDP-43 K181E KI affects cell-cell communications in differentiated organoids, we conducted CellChat analysis [39] using our single-cell transcriptomic data. We first compared outgoing and incoming signals for different cell types between HM K181E mutant and WT organoids. Our analysis suggested that in WT organoids, while radial glia have high outgoing and incoming signals (Fig. 3A, left panel), Exci-neurons only display intermediate levels of outgoing and incoming signals. By contrast, in HM K181E mutant organoids, exci-neurons display a substantial increase in both outgoing and incoming signals (Fig. 3A, right panel). Pathway analyses identified several signaling processes, represented by the PTN-PTPRZ1, PTPR-NTRK3, and SLIT1-ROBO1 ligand-receptor pairs, as key contributors to this disruption in Exci-neurons (Fig. 3B, C). Notably, the upregulated incoming signaling in Exci-neurons are accompanied by elevated expression of both ligands and receptors (Fig. 3D), suggesting a self-stimulated autocrine regulation. Among the three ligand-receptor pairs, the PTPRS-NTRK3 signaling axis was previously implicated in synaptic development and neuronal stress responses [40]. Its activation may represent a compensatory mechanism that counteracts synaptic dysfunction and neuronal stress in mutant cells. However, prolonged activation of this pathway seems to inadvertently exacerbate synaptic instability to contribute to TDP-43-associated neurodegeneration, as suggested previously [41].

**Figure 3.**
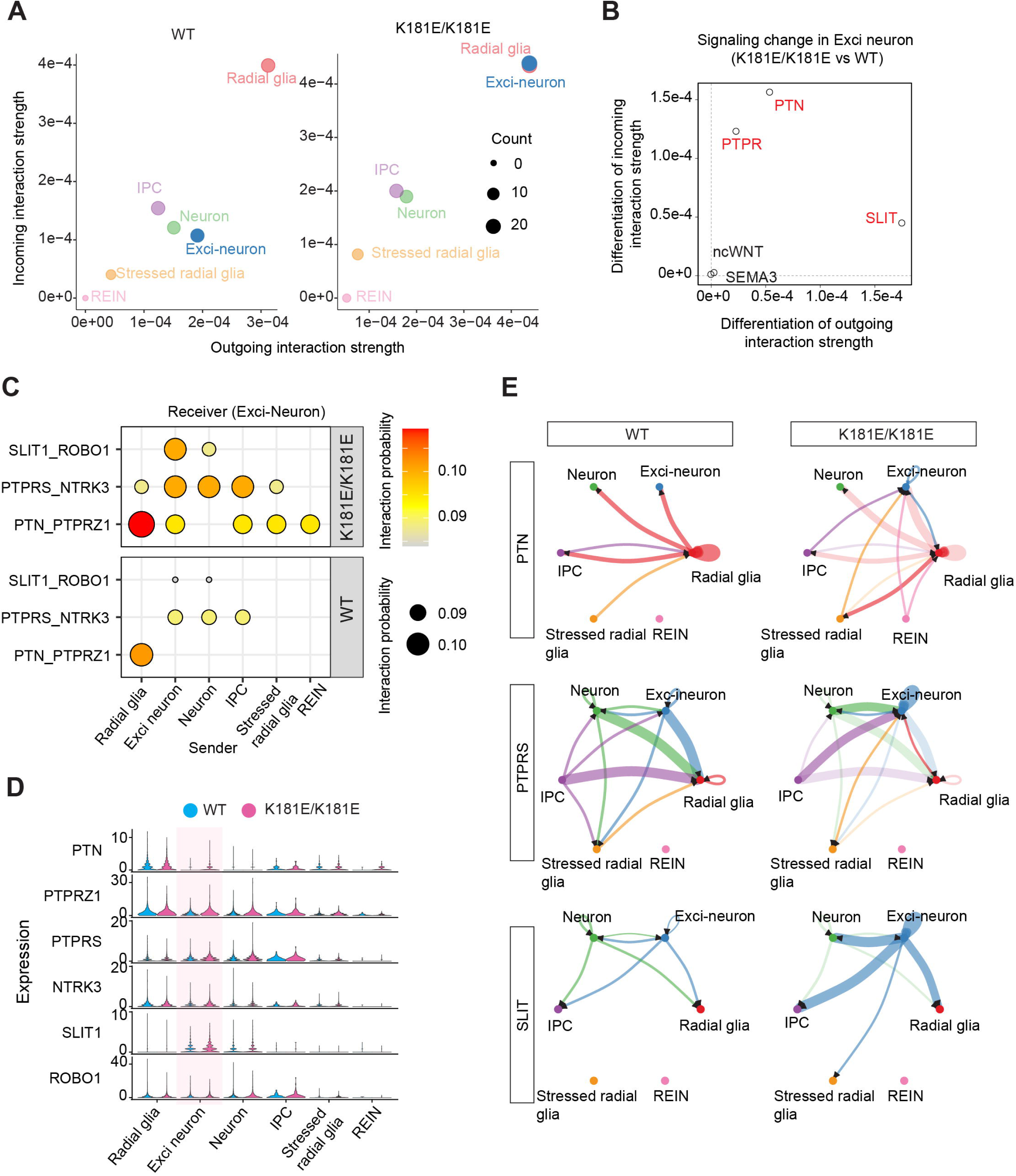
Single cell transcriptomic analysis reveals dysregulated cell-cell communications in TDP-43 K181E organoids. **(A)** Scatter plots showing the relative sender and receiver strength in WT (left) and K181E/K181E (right) forebrain organoids. x-axis and y-axis indicate the total outgoing or incoming interaction strength associated with each cell type. Dot size indicates the number of inferred ligand-receptor pairs associated with each cell type. Different cell types are color-coded. **(B)** Altered signaling dynamics in Exci-neurons in K181E/K181E organoids compared to WT organoids. Pathways significantly increased in K181E/K181E organoids were colored in red. **(C)** The relative contribution of the ligand-receptor pairs to the increased signaling activity in K181E/K181E organoids as shown in **(B)** between the indicated cell types as sender and Exci-neuron as receiver. The relative Interaction strength/probability is indicated by both color and dot size. **(D)** Violin plots showing the relative expression of the indicated genes in different cell types in WT and K181E/K181E forebrain organoids. Note that the most significant changes were observed in Exci-neurons (shaded). **(E)** Circle plots showing interaction strength of PTN (top), PTPR (middle), and SLIT (bottom) between different cell types in WT (left) and K181E/K181E (right) organoids. Node size and edge size indicate the corresponding interaction strength. Arrows indicate the signaling direction.

Additionally, we observed altered paracrine signals in these three pathways in HM K181E mutant organoids including de novo interactions via the PTN-PTPRZ1 axis between REINs, Radial glia and Exci-neurons (Fig. 3E). Immunostaining using PTPRZ1 antibodies confirmed the upregulation of PTPRZ1 expression in the forebrain layer of HM K181E organoids where Exci-neurons are enriched (Supplementary Fig. S4A, B). Since PTN and PTPRZ1 were known to activate the release of cytokines such as TNFα, IL6 and IL1β [42], these new interactions may promote neuroinflammation. Indeed, immunostaining with TNFα antibodies detected an upregulation of TNFα expression in HM K181E organoids (Supplementary Fig. S4C, D). Collectively, our findings reveal abnormal signaling networks and intercellular communications in mutant organoids that alter synaptic regulation and promote pro-inflammatory signaling responses.

### Altered RNA recognition and processing in TDP-43 K181E organoids

Since TDP-43 is an RNA binding protein, we analyzed RNA binding and processing by TDP-43 in WT and HM TDP-43 K181E organoids using eCLIP, a crosslinking and immunoprecipitation-based RNAseq assay for capturing global RNA-protein interactions (Fig. 4A, Supplementary Fig. S5A) [43]. The analysis identified 33,069 and 25,675 binding events for WT and K181E mutant, corresponding to 5,143 and 4,354 genes, respectively (Fig. 4B). Among these genes, 1,260 (22.4%) bound specifically to WT TDP-43, and 471 (∼8%) bound to TDP-43 K181E only. While the majority (69.2%) of the identified RNAs bound to both WT and mutant TDP-43, their affinity to these proteins was different, with 3016 RNAs showing reduced and 893 RNAs showing increased binding to the K181E mutant (Fig. 4C). GO pathway analysis on RNAs with reduced affinity to TDP-43 K181E revealed that TDP-43 binding deficiency predominantly occurred in genes involved in synapse assembly and trans-synaptic signaling (Fig. 4D) and mRNA metabolism (Supplementary Fig. S5B, Supplementary Table S3). Thus, the disease mutation alters TDP-43’s RNA-binding landscape, affecting primarily RNA metabolism and synaptic communications.

**Figure 4.**
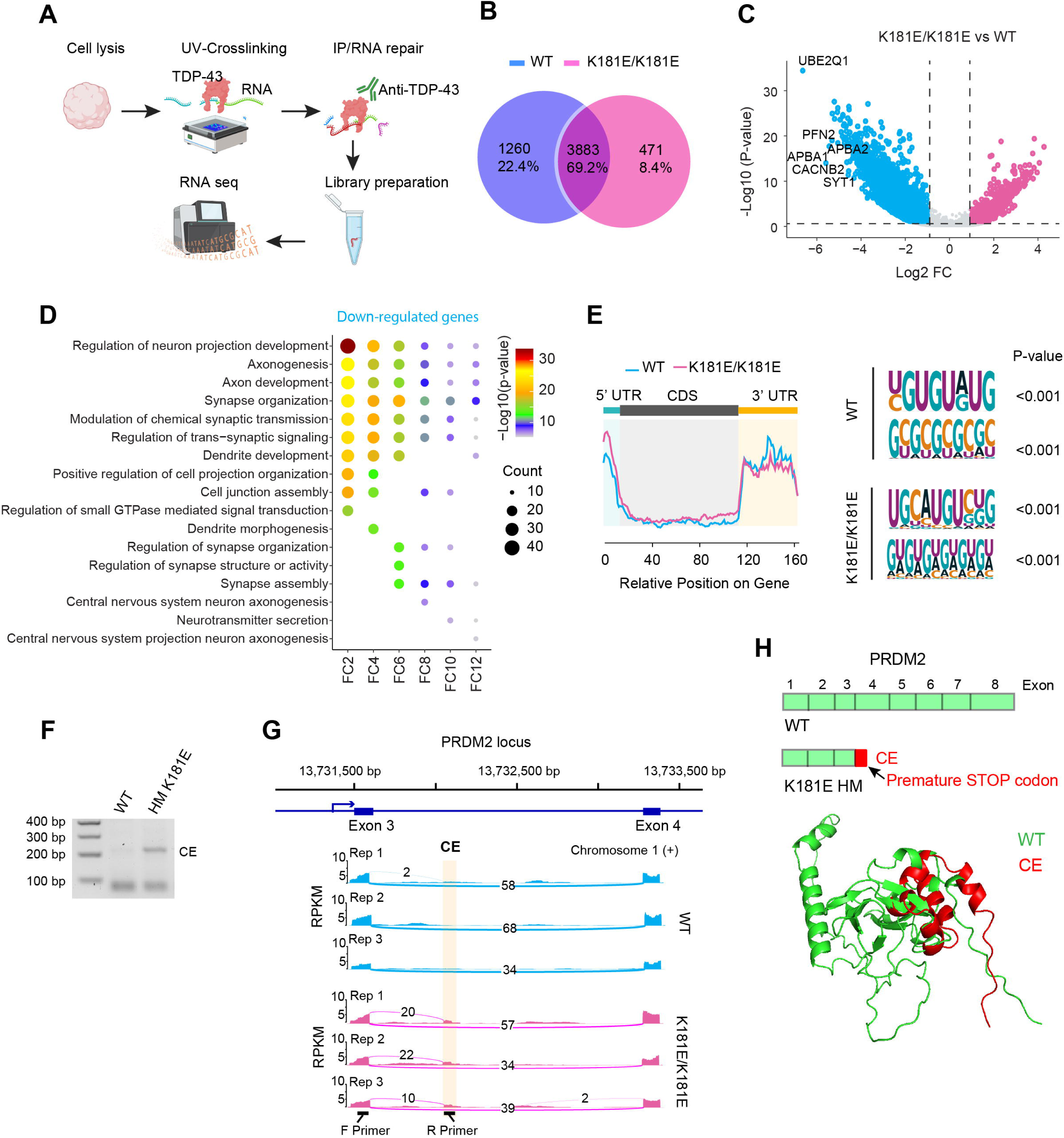
Altered RNA recognition and processing by the TDP-43 K181E mutant. **(A)** Schematic of the eCLIP workflow. **(B)** A Venn diagram summarizes TDP-43-associated RNAs detected by eCLIP in WT and TDP-43 K181E/K181E organoids. **(C)** Volcano plot showing RNAs differentially bound by TDP-43 in K181E/K181E organoids compared to WT organoids. P-value < 0.05 and foldchange (FC) ≥2 was used to identify differentially upregulated (red) or downregulated (blue) binding sites in TDP-43 K181E/K181E organoids compared to WT organoids. **(D)** Pathway analysis of RNAs with reduced TDP-43 binding in TDP-43 K181E mutants compared to WT organoids. Genes were grouped by the indicated fold-change (FC) of reduced binding before pathway analysis. Colors represent the -log_10_ (P-value) of pathway enrichment. **(E)** Metagene analysis (left panel) reveals average number of peaks enriched in WT and the K181E/K181E mutant across different RNA regions (Y-axis) and peak’s relative position on gene (X-axis). Detected TDP43 RNA recognition motifs in WT and K181E/K181E mutant (right panel), p < 0.001 by Yates’s Chi-Square test. **(F)** RT-PCR and gel electrophoresis confirms the presence of the predicted CE in the *PRDM2* transcript in K181E/K181E organoids. **(G)** Sashimi plot showing the inclusion of a cryptic exon (CE) in the *PRDM2* mRNA in K181E/K181E organoids in 3 replicates. **(H)** Schematic shows that the CE in *PRDM2* introduces an in-frame premature stop codon between exon 3 and exon 4 (top). The graph below shows an alpha-fold modeled structure of the translational product PRDM2* that contains the cryptic peptide (red) and its alignment with the structure of the corresponding WT PRDM2 N-terminal region.

Despite the differential RNA binding activity, sequence analysis showed that both WT and the K181E TDP-43 mutant bind predominantly to non-coding regions including introns and 5’- and 3’-UTRs, albeit with distinct sequence preference (Fig. 4E, Supplementary Table S4). This result suggests that the disease mutation disrupts TDP-43’s canonical RNA-binding specificity, causing a global change in RNA binding.

Since TDP-43 is known to regulate RNA splicing, particularly in suppressing cryptic exon (CE) inclusion [44–47], we investigated RNA splicing defects in HM TDP-43 K181E mutant organoids. To this end, we performed high reads RNAseq using bulk RNAs isolated from WT and K181E HM organoids and applied a well-established splicing analysis method named rMATS [48] to 500 million reads. This analysis identified 48 candidate RNA splicing events that might be altered in mutant organoids (Supplementary Table S5). Among them, 8 genes appeared to contain a CE in mutant organoids. As expected, we detected a cryptic splicing site in STMN2 (Supplementary Table S5), a well-established neuronal gene whose mis-splicing occurs in TDP-43-associated proteinopathies [49, 50]. To experimentally validate this result, we performed RT-PCR/qRT-PCR using primer pairs designed to specifically amplify CE-containing transcripts, with one primer located within the predicted CE sequence and one in a proximal exon. In addition to PRDM2, CE-containing products were detected for STMN2, ACTL6B, and SLC24A3 in HM K181E organoids (Fig. 4F and Supplementary Fig. S5G). These results support the conclusion that nuclear TDP-43 depletion in HM mutant organoids is accompanied by cryptic exon inclusion.

Among the genes analyzed, the most significant change occurred in PRDM2, which encodes an epigenetic factor regulating cellular senescence [51]. We therefore determined the transcript structure and predicted coding consequence of mis-spliced PRDM2. In HM K181E mutant organoids, ∼14% PRDM2 mRNAs has a CE of 490 bp long, derived from an intron sequence between exon 3 and 4 (Fig. 4G). Sanger sequencing confirmed the identity of the PRDM2 CE-containing RT-PCR product (Supplementary Fig. S5C). The mis-spliced *PRDM2* RNA bears an in-frame premature stop codon (Fig. 4H). Upon translation, it generates a truncated PRDM2 mutant (PRDM2*) of 56 amino acids, containing a 14 amino acid C-terminal cryptic peptide (NH_2_-VPDTLWTLSRYLNE-COOH) translated from the CE (Supplementary Fig. S5D, Fig. 4H). Although mass spectrometry did not detect this cryptic peptide in mutant organoids likely due to its low abundance and/or instability, it revealed ∼15% reduction in total PRDM2 protein level in mutant organoids after 135 days in culture (Supplementary Fig. S5E). The comparable PRDM2 CE inclusion and PRDM2 protein reduction suggest that CE inclusion reduces PRDM2 translational output, possibly through transcript or protein instability.

Collectively, these results demonstrate that the K181E mutation alters the TDP-43 RNA-binding specificity, causing a major defect in RNA splicing.

### PRDM2 CE-peptide immunoreactivity is detected in spinal motor neurons of ALS patients

To confirm PRDM2 mis-splicing at protein levels, we developed an antibody against the predicted CE-derived peptide. To validate the specificity of affinity-purified PRDM2 CE antibodies, we expressed mCherry-tagged WT PRDM2 or PRDM2 CE peptide fusion protein in iPSC-derived motor neurons and performed immunostaining with the PRDM2 CE-peptide antibody. Under identical staining and imaging conditions, CE-peptide immunoreactivity was detected only in neurons expressing PRDM2-CE-mCherry, but not in neurons expressing WT PRDM2-mCherry or mCherry, supporting the specificity of this antibody (Supplementary Fig. S5H). Interestingly, compared to nucleus-enriched WT PRDM2, the presence of CE caused more PRDM2 to accumulate in the cytoplasm.

Having validated the antibody specificity in neuronal cells, and because TDP-43-mediated RNA mis-splicing a shared molecular pathology across vulnerable neuronal populations in both FTD and ALS [9, 45, 52], we examined PRDM2 CE-peptide immunoreactivity directly in postmortem spinal cord tissues from ALS patients. To this end, we examined ventral horn spinal cord sections from ALS cases and age-matched controls and stained for ChAT together with an affinity-purified antibody raised against the PRDM2 CE–encoded peptide. Under identical acquisition settings, PRDM2 CE–peptide immunoreactivity was minimal in control ChAT+ motor neurons, whereas a subset of ChAT+ motor neurons in ALS sections exhibited detectable CE–peptide immunoreactivity, frequently with a cytosolic distribution. Quantification was performed on multiple sections per donor by investigators blinded to diagnosis, with sections treated as technical replicates nested within donors (Supplementary Fig. S5I, J).

To determine whether PRDM2-CE peptide immunoreactivity was associated with TDP-43 pathology, we performed additional co-immunostaining for p-TDP43 (Ser 409/410) and PRDM2-CE peptide in healthy control and ALS spinal cord sections. Because ChAT co-staining was not compatible with this antibody combination, this analysis was restricted to large ventral horn neuron-like cells identified by anatomical location and morphology. Compared with control tissue, ALS sections showed increased p-TDP-43 and PRDM2 CE-peptide signals, co-localizing in the cytoplasm. Importantly, PRDM2 CE-peptide immunoreactivity was predominantly observed in cells with elevated p-TDP-43 signal (Fig. 5A, B). These results confirm PRDM2 mis-splicing and CE inclusion in ALS patients and further link it to TDP-43 pathology.

**Figure 5.**
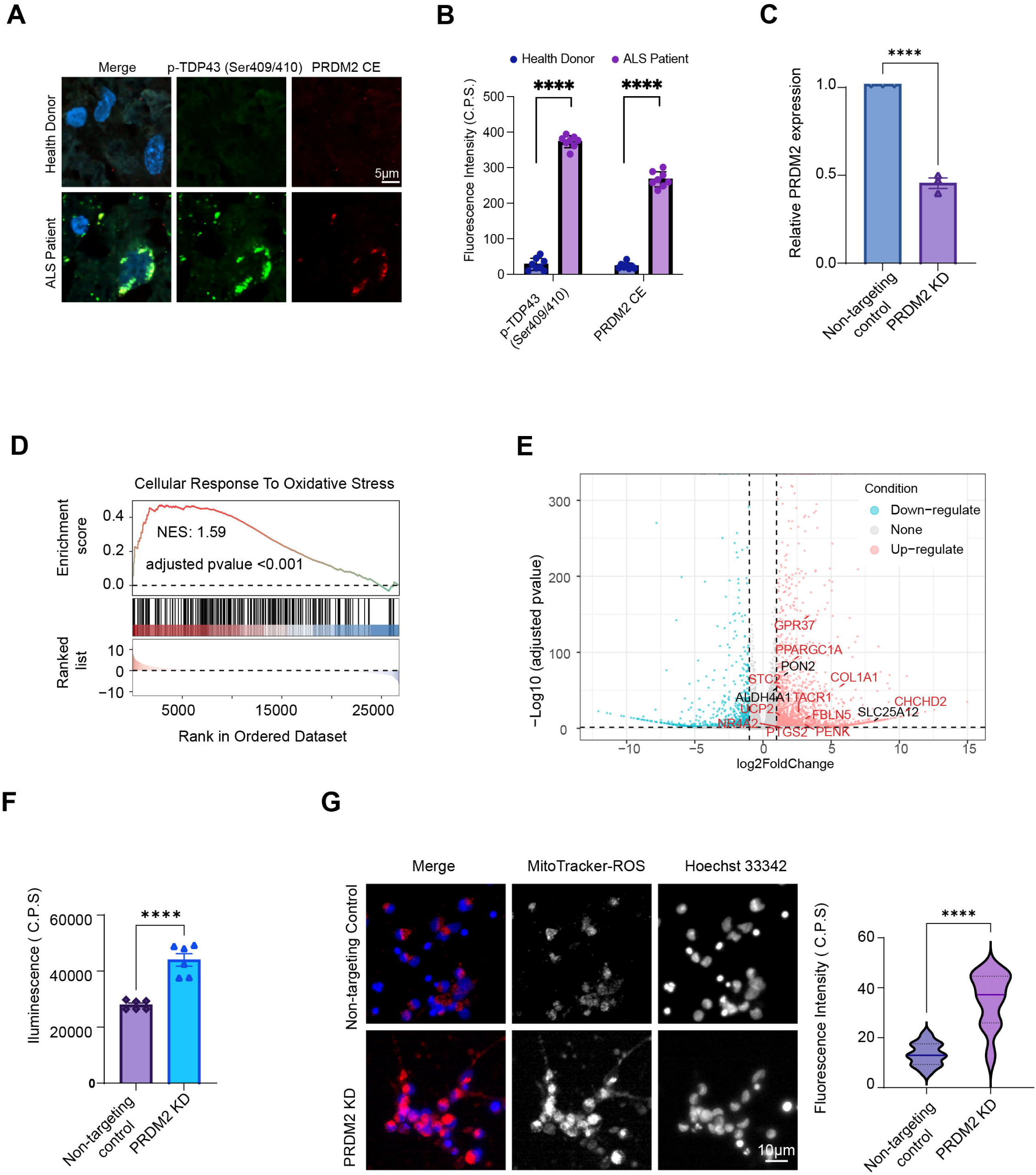
PRDM2 cryptic exon inclusion in ALS spinal motor neurons and PRDM2 knockdown-induced mitochondrial oxidative stress. **(A)** Representative confocal images of postmortem spinal cord sections from healthy controls and ALS patients co-stained with antibodies recognizing phospho-TDP-43 Ser409/410 and PRDM2 CE-peptide. Nuclei stained with DAPI. PRDM2 CE-peptide immunoreactivity was minimal in healthy control sections but was detected in motor neurons with elevated p-TDP-43 signal in ALS tissue (ventral horn region); ChAT not included due to compatibility; cell identity was inferred from anatomical location and morphology. Scale bar, 5 μm. **(B)** Quantification of p-TDP-43 Ser409/410 and PRDM2 CE-peptide fluorescence intensity as shown in (A). Data were obtained from sections derived from two healthy donors and two ALS patients. Each data point represents one analyzed cell. Error bars indicate mean ± s.e.m.; ****P < 0.0001 by unpaired Student’s t-test. **(C)** qPCR quantification of PRDM2 knockdown efficiency in motor neuron realtive to non-targeting control. Each data point represents an independent sample. Error bars indicate mean ± s.e.m. ****P < 0.0001, unpaired Student’s t-test, N=3. **(D)** GSEA of bulk RNA-seq data from PRDM2 knockdown motor neurons showing enrichment of oxidative stress response gene sets. NES, nominal P values, and adjusted P values are indicated. **(E)** Volcano plot of differential expressed genes in PRDM2 KD samples when compared to non-targeting control. **(F)** Extracellular H₂O₂ levels in conditioned medium measured by ROS-Glo H₂O₂ assay. Luminescence is shown as counts per second. Data are mean ± SEM; unpaired Student’s t test; ****P<0.0001; N=3. **(G)** Representative live-cell images and quantification of MitoTracker Red CM-H₂XRos fluorescence in control and PRDM2 knockdown motor neurons; Hoechst 33342 labels nuclei. Each point represents one motor neuron. Data are mean ± SEM; unpaired Student’s t test; ****P<0.0001; N=3. Scale bar, 10 μm.

### PRDM2 knockdown induces mitochondrial oxidative stress in motor neurons

Although PRDM2 CE inclusion in K181E/K181E organoids was associated with a modest ∼15% reduction in PRDM2 mRNA and protein expression, analysis of FTLD-TDP patient transcriptomic data revealed a more pronounced reduction (∼50%) in PRDM2 mRNA expression (Supplementary Fig. S5E-F), suggesting that PRDM2 reduction may be more accumulative in human disease contexts. We therefore used siRNA-mediated PRDM2 knockdown in terminally differentiated human motor neurons to evaluate the functional consequence of PRDM2 depletion in neurons. PRDM2 knockdown efficiency was confirmed by qRT-PCR (Fig. 5C).

Since PRDM2 was reported to function as transcriptional regulator, we characterized the transcriptional changes associated with PRDM2 depletion by bulk RNA sequencing. Differential expression analysis identified 2,312 upregulated (log2 FC >1, P-value<0.05) and 1,299 downregulated genes (log2 FC< -1, P-value<0.05) in PRDM2-depleted motor neurons when compared to control cells. Pathway analysis revealed cell response to oxidative stress-related genes among the most upregulated pathways (Fig. 5D). Notably, enrichment of this pathway was also observed in TDP-43 K181E homozygous forebrain organoids (Supplementary Fig.S6A) and in transcriptomic profiles from FTD patient data (Supplementary Fig.S6C), supporting the disease relevance of this transcriptional response.

In parallel, PRDM2 KD was associated with coordinated upregulation of oxidative stress-related genes in mitochondria (Fig. 5D). For example, among the most strongly induced transcripts was SLC25A12 (log2 FC=8.1), a mitochondria aspartate-glutamate carrier that supports malate-aspartate shuttle-mediated redox transfer and regulates NADH flux between the cytosol and mitochondria. ALDH4A1, a mitochondrial enzyme in proline metabolism that generates NADH, was also upregulated, consistent with increased delivery of reducing equivalents to the respiratory chain and a metabolic state that can favor mitochondrial ROS production under stress [53, 54]. In addition, the mitochondrial biogenesis and metabolic regulators PPARGC1A (log2 FC=2.27) and PPARGC1B (log2 FC=3.13) were also significantly upregulated, together with core genes in cellular response to oxidative stress (Fig.5E), These changes include PON2 (log2 FC=1.48), an intracellular antioxidant enzyme that limits mitochondrial superoxide production[55] and UCP2 (log2 FC=1.28), a mitochondrial uncoupling protein that reduces electron leakage and ROS generation by lowering membrane potential(Fig. 5E)[56] [57].

In addition to oxidative stress related pathways, enrichment of extracellular matrix organization and cell adhesion pathways was also detected following PRDM2 depletion. Importantly, similar enrichment of extracellular matrix–related transcriptional programs was observed in TDP-43 K181E homozygous organoids and in patient-derived transcriptomic datasets (Supplementary Fig.S6A–C, table S7). The overlap of oxidative stress–associated and extracellular matrix–associated transcriptional signatures across PRDM2-reduced motor neurons, TDP-43 mutant organoids, and patient tissue suggests that PRDM2 reduction is linked to a broader stress-responsive transcriptional state that recapitulates key molecular features of TDP-43–mediated pathology.

To assess oxidative stress at the functional level, we measured hydrogen peroxide levels in conditioned medium using the ROS-Glo™ H₂O₂ assay. PRDM2 knockdown cultures showed a significant increase in extracellular hydrogen peroxide compared to controls (Fig. 5F). Consistent with this result, live-cell staining with MitoTracker Red CM-H₂XRos detected increased mitochondrial reactive oxygen species in PRDM2-depleted motor neurons compared to controls (Fig. 5G).

## Discussion

Our study defines the molecular and cellular consequences of ALS- and FTD-associated TDP-43 K181E mutation under endogenous expression conditions and provides insight into how altered TDP-43 function drives neurodegeneration. By examining a human iPSC-derived neuronal system carrying the K181E mutation, we show that this mutation induces spontaneous TDP-43 hyperphosphorylation, cytoplasmic mislocalization, RNA dysregulation, neuroinflammatory signaling, neuronal stress and apoptosis without TDP-43 overexpression or exogenous stress. The close overlap between the transcriptional changes in K181E mutant cultures and those reported in FTD patient tissues further supports the disease relevance of the model established by this study. However, since most TARDBP mutations identified in ALS and FTD patients are heterozygous, the K181E/K181E organoid model should not be interpreted as a direct genetic replica of the patient condition, but rather a sensitized human cellular system that enables robust detection of cellular and molecular deficiencies associated with TDP-43 dysfunction.

The relatively young neural populations in forebrain organoids also fail to recapitulate aging-associated aspects of ALS and FTD. However, since molecular defects (e.g. cytoplasmic p-TDP-43 accumulation, nuclear TDP-43 depletion, and cryptic exon inclusion) revealed by the K181E organoid system arise from loss of endogenous TDP-43 function, these phenotypes should be broadly associated with ALS and FTD including cases bearing no TDP-43 mutations. Future studies incorporating aging-associated disease modulators are needed to determine how these early TDP-43 dependent RNA-processing defects evolve during disease progression.

A key advantage of the experimental framework used here is the ability to examine TDP-43 dysfunction within a multicellular neural context that supports endogenous cell–cell communications. This enabled identification of altered intercellular signaling networks involving SLIT1–ROBO1, PTN–PTPRZ1, and PTPR–NTRK3 ligand–receptor pairs, particularly within excitatory neurons. These signaling changes are consistent with disrupted synaptic regulation and enhanced neuroinflammatory responses reported in ALS and FTD [40, 42, 58]. The observation that excitatory neurons exhibit markedly increased incoming and outgoing signaling highlights their heightened vulnerability to TDP-43 dysfunction and suggests that maladaptive autocrine and paracrine signaling may contribute to disease progression.

The reconstitution of aberrant intercellular signaling in 3D organoid underscores the complex interplays between distinct cell populations in disease progression. In this regard, restoring neuronal connectivity and reducing neuroinflammatory responses through targeted interventions may offer new therapeutic avenues. For instance, targeting the PTN-PTPRZ1 signaling axis, known to elicit pro-inflammatory responses, may provide additional benefit by attenuating neuroinflammation and preserving neuronal function.

Among the cell types identified in organoids, we observed a mutant-enriched cell population characterized by elevated expression of ribosomal and translation-related genes, which we termed REIN. Hierarchical clustering based on cell type-averaged transcriptional profile suggest REINs have a neuronal identity. However, because organoids represent a developing neuronal system, the relatively immature property of REINs could reflect a mutation-associated differentiation bias, an early stress-associated neuronal state, or both. Additionally, although REIN-associated gene signatures are elevated in subsets of ALS and FTLD patient cells, this is not strictly neuron-specific. Thus, REINs should be interpreted as a mutant-enriched cell population in a neuronal transcriptional state and with altered translation-related gene expression.

Another key finding is the identification of widespread RNA dysregulation driven by the TDP-43 K181E mutation. A previous study reported that this TDP-43 mutant fails to bind a canonical TDP-43-binding motif *in vitro* [19]. Our unbiased eCLIP analysis reveals that the K181E mutation alters the TDP-43’s RNA-binding specificity, particularly in introns and 5’- and 3′-UTRs. Accordingly, we discovered several mis-splicing events in HM TDP-43 K181E organoids including CE inclusion in *PRDM2*. We also detected PRDM2 CE–peptide immunoreactivity in ALS spinal motor neurons, supporting the disease relevance of PRDM2 mis-splicing–associated pathology. Mechanistically, our eCLIP data did not reveal altered TDP-43 binding immediately adjacent to the CE splice junctions of PRDM2. Thus, the current data do not establish a direct splice-site–proximal mechanism for PRDM2 CE regulation. Nevertheless, our finding that CE inclusion is a major splicing defect in TDP-43-associated proteinopathies corroborate recent reports [45, 47], underscoring the clinical relevance of our model.

CE inclusion in *PRDM2* introduces a premature stop codon known to trigger non-sense-mediated RNA decay (NMD). Indeed, the *PRDM2* mRNA is downregulated significantly in FTD patients. However, the magnitude of PRDM2 mRNA reduction in K181E/K181E organoids was modest, which is consistent with the interpretation that the organoid system captures early molecular consequences of TDP-43 dysfunction rather than the cumulative changes that occur during the full course of disease progression.

CE inclusion usually results in altered translational products with reduced function or a gain-of-toxic function [45]. The change in TDP-43 binding to 5’- and 3’-UTRs may also affect protein translation given the established role of TDP-43 in translational regulation [59]. For *PRDM2*, the CE-containing mRNA encodes a truncated protein with a 14 amino acid carboxyl cryptic peptide derived from the CE. This and other abnormal translation products may have dominant gain-of-function activities contributing to the dose effect of the K181E allele on TDP-43 proteinopathies. Future work is needed to determine the abundance of cryptic peptide-containing proteins in FTD and ALS patients, which may serve as biomarkers for disease diagnosis and therapeutic evaluation.

An important implication of our findings is that PRDM2 mis-splicing downstream of TDP-43 dysfunction has functional consequences that extend beyond RNA dysregulation. PRDM2 is a chromatin-associated regulatory protein that has been reported to function as both a transcriptional repressor and a transcriptional activator through interactions with epigenetic coregulators and transcriptional machinery. Prior work supports a functional role for neuronal PRDM2 in maintaining stable transcriptional programs and neuronal circuit output [60]. A separate studyindicated that reduced neuronal PRDM2 is sufficient to shift neuronal state and behavioral output in vivo [61]. We modeled the loss-of-function outcome of PRDM2 mis-splicing using targeted knockdown in terminally differentiated motor neurons. This approach revealed that reduced PRDM2 expression induces mitochondrial oxidative stress, as evidenced by increased extracellular hydrogen peroxide and elevated mitochondrial reactive oxygen species, together with the activation of compensatory transcriptional programs involving mitochondrial redox flux, biogenesis, and uncoupling. In parallel, PRDM2 depletion triggered robust induction of extracellular matrix and adhesion programs intrinsic to motor neurons, accompanied by pronounced cell clustering and altered cellular architecture. Oxidative stress and extracellular matrix remodeling are both well-established features of ALS and FTD pathology, where mitochondrial dysfunction and altered cell–matrix interactions contribute to neuronal vulnerability [62–65]. Together, our findings suggest that PRDM2 reduction provides a plausible molecular link between TDP-43-dependent RNA mis-splicing and downstream metabolic and structural stress responses in motor neurons that are relevant to TDP-43–associated neurodegeneration (Supplementary Fig.S7). However, it should be noted that abnormal translation products from other mis-spliced mRNAs likely also contribute to the ALS/FTD disease phenotypes.

## Materials and methods

### Stem cell gene editing with knock-in K181E patient mutation

1.2×10^6^ Kolf2.1 NGN2 cells were nucleofected using Lonza P3 solution and program CA-137 with RNP complexes of 61 pmoles each of Alt-R™ S.p. HiFi and dCas9 Nucleases from IDT and 175 pmoles Synthego sgRNA targeting GACGCAAGTACCTTAGAATT plus 120 pmoles of IDT Ultramer TARDBP K181E editing oligo, 5’-AGCGACATATGATAGATGGACGATGGTGTGACTGCAAACTTCCTAATTCTGAGGTACTTGCGTCTGTGCTTTGGGAATTTTTGCCAACAAACTTCCTTAG-3’. Cells were plated on hrLaminin-521 coated plates in StemFlex medium supplemented with 1x RevitaCell and 1 µM IDT HDR Enhancer V2 and incubated at 32 °C. Medium was changed to StemFlex supplemented with 0.5X RevitaCell after 24 hours and to StemFlex after 48 hours. On the third day, single cells were sorted with a BD FACSMelody cell sorter at 1 cell per well to a Corning® Matrigel® Matrix coated 96-well plate with StemFlex medium supplemented with STEMCELL Technologies CloneR™2. Incubation temperature was shifted back to 37 °C and medium shifted to StemFlex by replacing 1/2 volume every other day. gDNA was isolated from expanded clones using DNAdvance on a Biomek4000. The target region was PCR amplified with 5’-TGGGAATGGAGTGTGTGAGT-3’ and 5’-TTCTCTTTTCCATCCATAGCGA-3’ primers using Phusion® Hot Start Flex DNA Polymerase. Product was purified with 0.8X AMPure XP beads and sequenced to identify both homozygous and heterozygous clones.

### Differentiation of iPSCs into forebrain neurons

The procedure was reported previously [66]. Briefly, the procedure for inducing neuronal differentiation of iPSCs using an induction medium (IM-N2) and a neuronal culture medium (CM). IM-N2 is prepared with Knockout DMEM/F12, N2 supplement, NEAA, Gluta-MAX, Chroman I, and doxycycline. On Day 0, iPSCs were observed for confluency, dissociated with Accutase, centrifuged, and resuspended in IM-N2 with Chroman I, and then seeded into Matrigel-coated plates. Over the next four days, cells were monitored microscopically for neurite extensions while media containing doxycycline is refreshed daily. By Day 4, cells exhibit neurite growth and are ready for replating onto poly-L-ornithine (PLO)-coated dishes, prepared in advance by coating with PLO solution, incubating, washing, and drying. Replated cells were cultured in CM comprising BrainPhys medium, B27+ supplement, neurotrophic factors (GDNF, BDNF, NT-3), laminin, and doxycycline, to promote neuronal maturation.

### Differentiation of iPSCs into forebrain organoids

3D culture and organoid growth were performed as previously described [67]. In brief, iPSCs grown on matrigel were dissociated into single cell suspension by Versene solution (ThermoFisher) and seeded into a 12-well Aggrewell plate (Stemcell Technologies) at 4,000 cells/ well. Next day, spheroids were transferred to an ultralow attachment plate (Corning) containing phase I medium: DMEM/F12, 20% Gibco KnockOut Serum Replacement, 1X Glutamax, 1X MEM Nonessential Amino Acid, 55 µM β-Mercaptoethanol, 1X Pen/Strep, 2 µM Dorsomorphine, 2 µM A83-01. After 5 days, medium was switched to phase II medium: DMEM/F12, 1X N-2 Supplement, 1X Glutamax, 1X MEM Nonessential Amino Acid solution, 1X Pen/Strep, 4 ng/mL WNT3a, 1 µM CHIR-99021, 1 µM SB-431542. On day 7, spheroids were embedded into matrigel and allowed to continue growing for 7 more days. On day 14, individual spheroids were manually freed from the matrigel and transferred to a SpinOmega bioreactor spinning at 120 RMP with phase III media: DMEM/F12, 1X N-2 Supplement, 1X B-27 Supplement, 1X Glutamax, 1X MEM Nonessential Amino Acid, 55 µM 2-Mercaptoethanol, 1X Pen/Strep, 2.5 µg/mL insulin. 50 days later, the medium was switched to final differentiation medium: Neurobasal medium, 1X B-27 Supplement, 1X Glutamax, 55 µM 2-Mercaptoethanol, 1X Pen/Strep, 0.2 mM Ascorbic acid, 0.5 mM cAMP, 20 ng/mL brain-derived neurotrophic factor (BDNF), 20 ng/mL glial-derived neurotrophic factor (GDNF). 107-day-old organoids were dissociated into single cells using a 50:50 mixed of Accutase and 0.25% Trypsin with DNase I (1 mg/mL) and plated onto chambered glass slides pretreated with 1% Matrigel (Corning, Inc). Alternatively, organoids were fixed and sectioned for immunostaining (see below).

### Tissue preparation, immunoblotting, and immunostaining

Organoids were fixed in 4% paraformaldehyde for 1 hour at room temperature, washed with PBS, and immersed in 15% sucrose/PBS solution overnight. Subsequently, organoids were embedded in O.C.T. compound (Sakura) in a plastic mold and frozen down in an ultralow freezer. Embedded organoids were sliced onto glass slides using a Cryostat (Leica). Slides were rinsed with PBS, permeabilized with 0.5% Triton-X/PBS solution for 1 hour at room temperature and blocked using 1% donkey serum in 0.1% Tween-20/PBS solution for 1 hour. Primary antibodies, chicken anti-TUJ1 (Abcam, ab41489,1/1000 dilution), rabbit anti-TDP-43 (10782-2-AP, 1/1000 dilution), mouse anti-phospho-TDP-43 (Cosmobio, CAC-TIP-PTD-M01A, 1/500 dilution), rabbit anti-cleaved caspase 3 (Cell signaling, 9664, 1/1000 dilution), were added to the slides and the slides were incubated in a humidified chamber at 4 °C overnight. After several washes in 0.1% Tween-20/PBS solution, DAPI (Sigma) or secondary antibodies, donkey anti-rabbit, anti-mouse, and anti-chicken (Jackson ImmunoResearch), diluted 1:1000 in blocking solution, were added to the slides. After 1 hour of incubation at room temperature, the slides were washed and mounted in antifade mounting solution (Fisher scientific).

For immunoblotting analyses, cells or organoids were first solubilized in NP40 lysis buffer containing 25 mM of Tris-HCl, pH 7.4, 150 mM of NaCl, 2 mM of MgCl_2_, 2 mM of KCl, and 0.5% of NP40, 1mM DTT and a protease inhibitor cocktail. After centrifugation at 20,000 g for 5 min, cleared supernatants were analyzed by immunoblotting with TDP-43 antibody (Proteintech, 10782-2-AP) and phospo-TDP-43 antibody (Cosmobio, TIP-PTG-M01A) at 1:500 dilution.

### Human spinal cord tissue and immunofluorescence staining

Postmortem human spinal cord tissues were obtained from two neurologically normal healthy controls and three patients with TDP-43 pathology and a genetic diagnosis of familial amyotrophic lateral sclerosis type4 (ALS4)[68]. All tissue samples were collected and handled in accordance with institutional guidelines and ethical approvals. Spinal cord segments containing the ventral horn were fixed in 4% paraformaldehyde, cryoprotected, and sectioned at 30 μm thickness for free-floating immunofluorescence staining. For immunostaining, all buffers were prepared in advance, including 1× Tris-buffered saline (TBS), TBS-A (TBS supplemented with 0.1% Triton X-100), and TBS-B (TBS-A supplemented with 5% bovine serum albumin). Blocking solution consisted of TBS-B supplemented with 5% normal horse serum. On day 1, sections were washed twice in TBS for 5 min each at room temperature. Antigen retrieval was not performed. Sections were then incubated sequentially in TBS-A for 15 min and TBS-B for 30 min, followed by blocking in blocking solution for 1 h at room temperature.After blocking, sections were incubated overnight at 4 °C with gentle agitation in primary antibody solution prepared in blocking buffer. Primary antibodies included antibodies recognizing the PRDM2 cryptic exon–specific sequence (Customized order from Thermofisher Scientific) and choline acetyltransferase (ChAT) to label spinal motor neurons. On day 2, sections were transferred to netwells and washed twice in TBS-A for 5 min each, followed by a 15 min wash in TBS-B. Sections were then incubated for 1 h at room temperature in the dark with species-appropriate Alexa Fluor–conjugated secondary antibodies diluted in blocking solution. After secondary antibody incubation, sections were washed three times in TBS.

To reduce tissue autofluorescence, sections were treated with freshly prepared 1× TrueBlack solution diluted in 70% ethanol for 4 min, followed by three washes in TBS. Sections were mounted onto glass slides and coverslipped using DAPI Fluoromount to counterstain nuclei. Slides were allowed to dry overnight in the dark, and coverslips were sealed with nail polish prior to imaging. Fluorescence images were acquired using a confocal microscope under identical acquisition settings for all conditions. Quantification of PRDM2 cryptic exon–positive motor neurons was performed on multiple sections per sample by investigators blinded to sample identity.

### Lentiviral overexpression and validation of PRDM2 cryptic exon peptide antibody specificity

To validate the specificity of the PRDM2 cryptic exon peptide antibody, lentiviral vectors were generated to express mCherry-tagged WT PRDM2, mCherry-tagged PRDM2 containing the cryptic exon-encoded peptide sequence (CP), or mCherry alone. Transgene expression was driven by the neuronal synapsin promoter to restrict expression to neurons. Lentiviral particles were produced using standard lentiviral packaging procedures and used to infect iPSC-derived motor neurons. Motor neurons were purchased from BrainXell and cultured according to manufacture’s protocol. After 7 days post-plating, neurons were infected with the indicated lentiviruses and maintained for 48 h to allow transgene expression. Cells were then fixed with 4% paraformaldehyde, washed with PBS, permeabilized, and blocked using the same immunofluorescence staining procedure described above. Cultures were incubated with antibodies recognizing TUJ1 and the affinity-purified PRDM2 cryptic exon peptide antibody, followed by species-appropriate fluorescent secondary antibodies. mCherry fluorescence was used to identify infected cells expressing WT PRDM2-mCherry, PRDM2-CE-mCherry, or mCherry control. Images were acquired under identical imaging settings across conditions. Antibody specificity was assessed by comparing PRDM2 CE-peptide immunoreactivity among mCherry-positive neurons expressing WT PRDM2-mCherry, PRDM2-CP-mCherry, or mCherry alone. Only PRDM2-CE-mCherry–expressing neurons showed detectable PRDM2 CE-peptide immunoreactivity under these conditions.

### siRNA-mediated PRDM2 knockdown in human iPSC-derived motor neurons

Human iPSC-derived spinal motor neurons were obtained from BrainXell and cultured according to the manufacturer’s protocol until terminal differentiation. Motor neurons were seeded onto poly-D-lysine– and laminin-coated plates and maintained in BrainXell motor neuron maintenance medium. All knockdown experiments were performed on mature motor neurons to avoid perturbing lineage specification or early differentiation. PRDM2 knockdown was achieved using Accell siRNA technology (Horizon Discovery), following BrainXell’s recommended gene silencing protocol for post-mitotic neurons . Accell siRNAs targeting human PRDM2 and a non-targeting negative control Accell siRNA were resuspended to 100 μM stock solutions using 1× siRNA buffer prepared from 5× siRNA buffer diluted in RNase-free water. siRNA stock solutions were mixed thoroughly and incubated at room temperature with gentle agitation to ensure complete dissolution before aliquoting and storage at −80 °C. For knockdown experiments, siRNA delivery was initiated five days after motor neuron seeding. On the day of transfection, culture medium was gently removed and replaced with Accell siRNA Delivery Medium containing either PRDM2-targeting Accell siRNA or non-targeting control siRNA at a final concentration of 300 nM. Cells were incubated in Accell delivery medium for 24 h at 37 °C. The following day, Accell delivery medium was completely removed and replaced with fresh motor neuron maintenance medium. Neurons were maintained for an additional 48 h prior to downstream analyses, resulting in a total knockdown duration of three days.

Knockdown efficiency was assessed at the mRNA level by RT–qPCR. Total RNA was extracted from motor neuron cultures using a column-based RNA isolation kit, followed by cDNA synthesis using a reverse transcription master mix according to the manufacturers’ instructions (Thermofisher Scientific). PRDM2 expression levels were normalized to housekeeping genes and compared with non-targeting control conditions. Functional assays, including measurements of extracellular hydrogen peroxide levels, mitochondrial reactive oxygen species, immunofluorescence imaging, and bulk RNA sequencing, were performed within this defined knockdown window to capture acute consequences of PRDM2 reduction in mature motor neurons.

### Image acquisition and processing

For analysis of p-TDP-43 accumulation in intact organoid sections, images were acquired from the outer organoid regions enriched for cortical neuron-like cells based on laminar organization. The same anatomical region was sampled across genotypes to enable comparison. Fluorescence confocal images were acquired using a Nikon CSU-W1 SoRa microscope equipped with temperature and CO_2_ control enclosures. Image reconstructions and analyses, including 3D and time lapse visualizations, were performed using Imaris software (licensed to NIH) or Image J. Fluorescence intensity was quantified using the open-source Fiji/Image J software. Images of randomly selected fields were split into individual channels. A consistent threshold method was applied to individual channel. The particle analysis function was used to automatically identify all other fluorescence structures for size and intensity measurement. Statistical analyses were conducted using Excel (for Student’s t-test) or GraphPad Prism versions 8.0 and 9.0. P-values were calculated using Student’s t-test in Excel or one-way ANOVA in GraphPad Prism. Curve fitting, including linear and nonlinear models, as well as IC_50_ calculations, were also performed using GraphPad Prism versions 8.0 and 9.0.

### Single-cell RNAseq cDNA library preparation

We mixed 30 organoids from multiple clones, two independent batches (15/batch) for scRNAseq. Single cells were captured and barcoded using the 10X Chromium platform (10X Genomics), and scRNA-seq libraries were prepared according to the Chromium Single Cell 3L Reagent Kits User Guide (v3). Gel Bead-In Emulsions (GEMs) were generated using single-cell preparations and processed with a 10X Chromium Controller. Following GEM reverse transcription and cleanup, mRNA from uniquely barcoded cells was converted into complementary DNA (cDNA), which was amplified to generate sufficient mass for library construction. The amplified cDNA underwent enzymatic fragmentation, size selection (targeting an insertion size of approximately 350 bp), end-repair, A-tailing, adaptor ligation, and dual-index PCR to construct the single-cell 3L gene expression libraries. Quality control was performed using Agilent Bioanalyzer High Sensitivity DNA chips, and libraries were pooled to ensure a similar number of reads from each single cell before sequencing on the NovaSeq S4 (Novogen Inc).

### Single-cell RNA-seq data processing

Reads from scRNA-seq were mapped to human transcriptome (GRCh38) using Cell Ranger software (10x Genomics) with the default parameters for each sample separately. Genes were retained with expressing in more than 5 cells. Cells with less than 200 detected genes were excluded. We also performed additional quality control steps including removing doublets (according to the multiplet rate provided by 10x Genomics based on the number of cells loaded and recovered) and filtering cells with the percentage of mitochondrial genes (≥ 10%). R package DoubletFinder (v2.0.4) was used to detect doublets with default parameters. A total of 16,549 cells passed the quality control in WT and K181E/K181E organoids. ‘SCTransform’ method from R package Seurat (v 5.1.0) was used to perform data normalization and scaling based on the top 2000 highly variable genes. The first 15 PCs were selected for tSNE and clustering analysis with default parameters after data reduction with PCA. Different subclusters of single cells were revealed from the tSNE plot.

### Cell type annotation

Differentially expressed genes (DEGs) for each cluster were first identified by ‘FindAllMarkers’ from Seurat (v 5.1.0) using the following criteria: adjusted p-value < 0.05, ≥10% of cells expressing the gene, and log₂ fold change ≥ 1 (Supplementary Table S1). These marker genes were analyzed using the ACT web server (http://biocc.hrbmu.edu.cn/ACT/), which employs a weighted and integrated gene set enrichment (WISE) method and knowledge-based resources for initial cell type characterization. The results were then manually refined based on the classical markers of different cell type, and the finalized cluster annotations with detailed gene marker expression are included in Supplementary Table S1.

### Construction of single cell trajectory

R package Monocle3 (v1.3.7) was used to learn the trajectory graph based on scRNA-seq data. Top 2000 most variable genes from ‘FindVariableFeatures’ and top 15 PCs were used to perform Uniform Manifold Approximation and Projection (UMAP) and Leiden clustering. The trajectory graph for single cells was learnt using R function (learn_graph) with default parameters. We utilized R package CytoTRACE2 (v1.0.0) to calculate the developmental potential score for single cell. The count matrix was set as input with default parameters.

### REIN score calculation

We used the expression of REIN signature genes to examine the enrichment differences of REIN phenotype in ALS/FTD patients [32]. REIN related genes were defined using R function “FindVariableFeatures” from Seurat (v5.1.0). Genes expressed in ≥ 20% of REIN with adjusted P-values < 0.05 and foldchange ≥ 4 were considered as REIN related genes (Supplementary Table S2). REIN score was calculated as the average expression of the identified REIN related genes.

Hierarchical clustering of annotated cell types

To evaluate transcriptional relationships among annotated cell types, normalized expression values were averaged for each cell type. Pairwise Pearson distances were calculated across cell type-averaged expression profiles, and hierarchical clustering was performed using the resulting distance matrix. The dendrogram was used to assess the relative transcriptional relationship between REINs and other annotated cell populations.

### Pathway enrichment analysis

Gene enriched biological pathways (Gene Ontology and Gene Set Enrichment Analysis) were detected using R package ClusterProfiler (v4.14.0) [69] with cutoff of P-value < 0.05. R package ggplot2 (v3.5.1) was used to plot the GO figures. The enriched KEGG pathways were obtained using Enrichr (https://maayanlab.cloud/Enrichr/).

### Cell-cell communication analysis

R package CellChat (v2.1.2) [39] was used to detect significant cell-cell communication signaling among different cell types. Secreted signaling in CellChat database was selected for downstream analysis. ‘trimean’ method was used to calculate communication probability of ligand-receptor signals. Ligand-receptor signals detected in at least ten cells were retained for downstream analysis.

### eCLIPseq library preparation and data analysis

eCLIP was performed by Eclipsebio according to the published single-end seCLIP protocol with the following modifications [43]. Approximately 40 mg of sample from each condition was cryogrinded and UV crosslinked at 400 mJoules/cm^2^ twice with 254 nm radiation. Cryoground tissue was then stored until use at -80 **°**C. Tissue aliquots were lysed using 750 μL of eCLIP lysis mix with a modified composition of 6 µL of a proteinase Inhibitor Cocktail and 20 µL of murine RNase inhibitor. Samples were then subjected to two rounds of sonication for 4 minutes with 30 second ON/OFF at 75% amplitude. A pre-validated anti-TDP-43 antibody (Eclipsebio) was then coupled to IgG Dynabeads (ThermoFisher), added to lysate, and incubated overnight at 4°C. Prior to immunoprecipitation, 2% of the sample was taken as the paired input sample, with the remainder magnetically separated and washed with eCLIP high stringency wash buffers. IP and input samples were cut from the membrane at the relative band size to 75kDa above. RNA adapter ligation, IP-western, reverse transcription, DNA adapter ligation, and PCR amplification were performed as previously described [43].

The eCLIP cDNA adapter contains a sequence of 10 random nucleotides at the 5’ end. This random sequence serves as a unique molecular identifier (UMI) after sequencing primers are ligated to the 3’ end of cDNA molecules. Therefore, eCLIP reads begin with the UMI and, in the first step of analysis, UMIs were pruned from read sequences using umi_tools (v0.5.1) [70]. UMI sequences were saved by incorporating them into the read names in the FASTQ files to be utilized in subsequent analysis steps. Next, 3’-adapters were trimmed from reads using cutadapt (v2.7), and reads shorter than 18 bp in length were removed. Reads were then mapped to a database of human repetitive elements and rRNA sequences compiled from Dfam [71] and Genbank. All non-repeat mapped reads were mapped to the human genome (hg38) using STAR (v2.6.0c) [72]. PCR duplicates were removed using umi_tools (v0.5.1) by utilizing UMI sequences from the read names and mapping positions. Peaks were identified within eCLIP samples using the peak caller CLIPper [73].

For each peak, IP versus input fold enrichment values were calculated as a ratio of counts of reads overlapping the peak region in the IP and the input samples (read counts in each sample were normalized against the total number of reads in the sample after PCR duplicate removal). A P-value was calculated for each peak by the Yates’ Chi-Square test, or Fisher Exact Test if the observed or expected read number was below 5. Comparison of different sample conditions was evaluated in the same manner as IP versus input enrichment; for each peak called in IP libraries of one sample type we calculated enrichment and P-values relative to normalized counts of reads overlapping these peaks in another sample type. Peaks were annotated using transcript information from GENCODE [74] with the following priority hierarchy to define the final annotation of overlapping features: protein coding transcript (CDS, UTRs, intron), followed by non-coding transcripts (exon, intron). HOMER de novo motif analysis pipeline was used to identify significantly enriched binding motifs of TDP-43 in each sample condition (eCLIPSEBIO) [75].

### Bulk RNA-seq library preparation and data analysis

800 ng of total RNA samples to generate the library with the NEBNext rRNA Depletion Kit v2 (Human/ mouse/rat) (NEB #E7405), NEBNext Ultra II Directional RNA Library Prep Kit for Illumina (NEB #E7760) and NEBNext^®^ Multiplex Oligos for Illumina^®^ (96 Unique Dual Index Primer Pairs, NEB, E6440) as described in the manufacturer’s manual. Sequencing carried out in the Illumina NextSeq2000 instrument with 101×101 pair end configuration by NCI genomic core.

Adapters were trimmed using Cutadapt (v4.7), and reads were aligned to the GRCh38 genome build with STAR (v2.7.11b) using gene annotations from GENCODE v42. Gene expression was quantified with HTSeq (v2.0.7) [76], utilizing the same gene annotations. For differential gene expression analysis, all samples were processed uniformly following the standard DESeq2 [77] workflow. Differential expressions were defined based on an adjusted P-value threshold of <0.05 and |log2 fold change (log_2_FC) | > 1. To identify gene expression rescued by exportin-1 inhibitor treatment, the fold change of expression level should be less than without inhibitor treatment, with P-value < 0.05.

### RNA Splicing Analysis

For differential splicing analysis, STAR-aligned BAM files from RNA-seq dataset were used as input for rMATS (v4.1.2) [48] with the GRCh38 reference genome. A ΔΨ (percent spliced in) threshold of 10% was applied to identify significant changes between groups. Junction files from rMATS were further analyzed using custom Python scripts to compute Ψ values for each splicing junction. ΔΨ values were calculated as the mean Ψ of three biological replicates. Cryptic splicing events were identified as splicing junctions that met the following criteria: Ψ < 5% in control samples, ΔΨ > 10%, and the junction was not annotated in GENCODE v42.

### RT-PCR validation of cryptic exon events

Selected cryptic exon events identified by RNA-seq/rMATS analysis were validated by RT-PCR using cDNA generated from WT and K181E/K181E organoid RNA. Primer pairs were designed to detect CE-containing transcripts, with one primer located within the predicted CE sequence and the paired primer located in an adjacent annotated exon. PCR products were resolved by agarose gel electrophoresis, and predicted CE-containing amplicons were identified based on expected molecular size. Because cryptic exon sequences are present as intronic genomic sequences in both WT and mutant cells, CE-directed primers may occasionally amplify low-abundance precursor RNA or nonspecific products. Therefore, only PCR products corresponding to the predicted CE-containing amplicon size were used for interpretation. Where indicated, PCR products were confirmed by Sanger sequencing.

### Whole exome sequencing and off-target analysis

10 ug of genomic DNA was extracted from WT stem cell and K181E/K181E stem cell, then sent out for whole exome sequencing provided by Azenta Life Sciences. Briefly, Sequencing adapters and low-quality bases in raw reads were trimmed using Trimmomatic 0.39. Cleaned reads were then aligned to the Homo sapiens GRCh38 reference genome using Sentieon 202308. After sorting the alignments, PCR and optical duplicates were flagged and removed. Single-nucleotide variants (SNVs) and small insertions and deletions (INDELs) were identified using Sentieon 202308 with the DNAscope algorithm. Variant call format (VCF) files generated by the pipeline were subsequently normalized with bcftools 1.13 by left-aligning INDELs and decomposing multiallelic sites into multiple records. Top 20 predicted off target sequences were obtained from CRISPOR and further cross-checked in the processed whole exome sequencing files (Table S6).

### RNA-seq data processing and differential expression analysis in PRDM2 KD motor neurons

RNA-seq reads were quality-controlled using FastQC and MultiQC and trimmed with fastp. Reads were aligned to the human GRCh38.p14 genome using STAR with GENCODE v47 annotations. Gene-level counts were generated using featureCounts. Differential expression analysis was performed in R using DESeq2 on raw count data with size-factor normalization and Wald tests, with FDR correction applied to p-values. Gene symbols were assigned using GENCODE v47 annotations. For visualization and quality control, variance-stabilized expression values (VST) were used to generate principal component analysis (PCA) plots and expression heatmaps. PCA plots were colored by experimental condition. Heatmaps were generated for the top differentially expressed genes (adjusted p-value < 0.05 and |log2 fold change| ≥ 1), ranked by adjusted p-value, with row-wise Z-score scaling applied to emphasize relative expression differences across samples. Gene symbols were assigned using GENCODE v47 annotations for display purposes.

### Protein structural simulation

WT PRDM2 and PRDM2+CE protein structures were simulated by AlphaFold Server(https://alphafoldserver.com/) and aligned by PyMOL.

### Statistic and reproducibility

All statistical analyses were conducted with GraphPad Prism v10 or Excel. Statistical methods and the number of cells or brain organoid samples (N) are indicated in figure legends or shown in figures as individual data point. Biological repeats (n) are specified in figure legends. We did not predetermine sample size. The sample sizes are consistent with similar studies reported in the literature. No biological repeat was excluded from the analyses. For individual cell analyses, a few data points (less than 2%) outside of 1.5 times the interquartile range were considered as outliers and were excluded. All experiments were repeated at least twice with individual data point labeled in figures unless specified. The immunoblotting is representative of similar results from at least two independent biological replicates unless specified in figure legends. For imaging analyses, cells in randomly selected fields were analyzed. The researchers were not blinded. All iPSC cell lines were derived from a ADRDs genetic risk-free clone, which was initially obtained from a male. Figures were prepared using ImageJ 1.54f, Imaris 9.9.0, Adobe Photoshop v25.12.1, and Adobe Illustrator 28.7.4.

### Sequencing Data availability

Raw and processed data files of scRNA-seq, bulk RNA-seq and eCLIP-seq can be accessed from Sequence Read Archive (SRA), NCBI with Bioproject ID: PRJNA1195353, http://www.ncbi.nlm.nih.gov/bioproject/1195353<u>. Code used in this project is available upon</u> <u>request.</u>

Whole exome sequencing files and processed sequencing data files can be found at: https://zenodo.org/records/14291940?preview=1&token=eyJhbGciOiJIUzUxMiJ9.eyJpZCI6IjFlOWM5MzU3LTQyMzUtNGQzYy05MTY2LTlhNDk4NzI5NWU5OSIsImRhdGEiOnt9LCJyYW5kb20iOiIwNDYzYWY0NjJhODExZWQ5YzkzNzI4MzlhYmZmNjc4ZiJ9.crjqSUlB8r6LPGyMV8YaB9zSLzSdO1zE33tjjF1UVXaQqnWpy8ZOW9FU6VPrbNN5zyf2JXJOpfDteT_RaaZ8Zg

## Abbreviations

ALS: Amyotrophic lateral sclerosis
ANOVA: Analysis of variance
BDNF: Brain-derived neurotrophic factor
CNS: Central Nervous system
CE: Cryptic exon
CM: Neuronal culture medium
CRISPR: Clustered regularly interspaced short palindromic repeats
cDNA: Complementary deoxyribonucleic acid
DAPI: 4’,6-diamidino-2-phenylindole
DEG: Differentially expressed gene
DMEM: Dulbecco’s Modified Eagle Medium
eCLIP: Enhanced crosslinking and immunoprecipitation
Exci-neuron: Excitatory neuron
FBS: Fetal bovine serum
FTD: Frontotemporal dementia
FTLD-TDP: Frontotemporal lobar degeneration with TDP-43 pathology
GDNF: Glial-derived neurotrophic factor
GO: Gene Ontology
HM: Homozygous mutant
HR: Homology-directed repair
iPSC: Induced pluripotent stem cell
IM-N2: Induction medium containing N2 supplement
IPC: Intermediate progenitor cell
KI: Knock-in
LCD: Low-complexity domain
LLPS: Liquid–liquid phase separation
MG132: Carbobenzoxy-Leu-Leu-leucinal (proteasome inhibitor)
NGN2: Neurogenin-2
NMD: Nonsense-mediated mRNA decay
PBS: Phosphate-buffered saline
PCA: Principal component analysis
PCR: Polymerase chain reaction
PLO: Poly-L-ornithine
PRDM2: PR domain zinc finger protein 2
PTN: Pleiotrophin
PTPRZ1: Receptor-type tyrosine-protein phosphatase zeta
qPCR: Quantitative polymerase chain reaction
RBP: RNA-binding protein
REIN: Ribo-enriched immature neuron
RIN: RNA integrity number
RNA-seq: RNA sequencing
RRM: RNA recognition motif
RT-PCR: Reverse transcription polymerase chain reaction
scRNA-seq: Single-cell RNA sequencing
TNFα: Tumor necrosis factor alpha
TDP-43: TAR DNA-binding protein 43
UMAP: Uniform manifold approximation and projection
UTR: Untranslated region
WT: Wild type

## Acknowledgements

We thank NCI CCR Genomics Core and the NIDDK Advanced Imaging Core with data acquisition. We thank K. Bharti (NIH/NEI) and W. Li (NIH/NEI), T. Zhang (NIH/NIA) for critical reading of the manuscript. We also thank the Therapeutic Development Branch Biology Group, Lara El Touny (NIH/NCATS), Dvir Blivis (NIH/NCATS), and Ty Voss (NIH/NCATS) for their valuable support.

## Author information

### Author contributions

Q.Z. conceived and supervised the study. Q.Z. and Y.Y. directed the experimental designs and interpreted the data. M.L. and X.F. performed computational analysis on single cell RNA seq and RNA splicing, respectively. J. Z. constructed the iPSC knock-in lines, N.C. performed 2D iNeuron-related experiments and analyses, Q.Z., X.Y., J.S., Y.C., S.A., S.L., J.V. C.G.and K.L. conducted other experiments and additional data analysis. Q.Z. and Y.Y. wrote the manuscript. All authors read, edited, and approved the manuscript.

### Corresponding authors

Correspondence to Qi Zhang, Wei Zheng or Yihong Ye.

### Data, code and materials availability

All data and code needed to evaluate and reproduce the results in the paper are present in the paper and/or the Supplemental Materials. This study did not generate new materials.

### Declarations

Ethics approval and consent to participate

Not applicable.

### Consent for publication

Not applicable.

## References

1. M. Neumann et al., Ubiquitinated TDP-43 in frontotemporal lobar degeneration and amyotrophic lateral sclerosis. Science 314, 130–133 (2006). 10.1126/science.1134108.

2. T. Arai et al., TDP-43 is a component of ubiquitin-positive tau-negative inclusions in frontotemporal lobar degeneration and amyotrophic lateral sclerosis. Biochem. Biophys. Res. Commun. 351, 602–611 (2006). 10.1016/j.bbrc.2006.10.093.

3. J. Brettschneider et al., Sequential distribution of pTDP-43 pathology in behavioral variant frontotemporal dementia (bvFTD). Acta Neuropathol. 127, 423–439 (2014). 10.1007/s00401-013-1238-y.

4. L. Guo, J. Shorter, Biology and pathobiology of TDP-43 and emergent therapeutic strategies. Cold Spring Harb. Perspect. Med. 7, a024554 (2017). 10.1101/cshperspect.a024554.

5. T. R. Suk, M. W. C. Rousseaux, The role of TDP-43 mislocalization in amyotrophic lateral sclerosis. Mol. Neurodegener. 15, 45 (2020). 10.1186/s13024-020-00397-1.

6. P. Tziortzouda, L. Van Den Bosch, F. Hirth, Triad of TDP43 control in neurodegeneration: Autoregulation, localization and aggregation. Nat. Rev. Neurosci. 22, 197–208 (2021). 10.1038/s41583-021-00431-1.

7. A. E. Conicella et al., ALS mutations disrupt phase separation mediated by α-helical structure in the TDP-43 low-complexity C-terminal domain. Structure 24, 1537–1549 (2016). 10.1016/j.str.2016.07.007.

8. F. Gasset-Rosa et al., Cytoplasmic TDP-43 de-mixing independent of stress granules drives inhibition of nuclear import, loss of nuclear TDP-43, and cell death. Neuron 102, 339–357.e7 (2019). 10.1016/j.neuron.2019.02.038.

9. J. P. Ling et al., TDP-43 repression of nonconserved cryptic exons is compromised in ALS-FTD. Science 349, 650–655 (2015). 10.1126/science.aab0983.

10. T. F. Gendron, L. Petrucelli, Rodent models of TDP-43 proteinopathy: Investigating the mechanisms of TDP-43-mediated neurodegeneration. J. Mol. Neurosci. 45, 486–499 (2011). 10.1007/s12031-011-9610-7.

11. T. Arai et al., Phosphorylated and cleaved TDP-43 in ALS, FTLD and other neurodegenerative disorders and in cellular models of TDP-43 proteinopathy. Neuropathology 30, 170–181 (2010). 10.1111/j.1440-1789.2009.01089.x.

12. L. M. Igaz et al., Dysregulation of the ALS-associated gene TDP-43 leads to neuronal death and degeneration in mice. J. Clin. Invest. 121, 726–738 (2011). 10.1172/JCI44867.

13. V. Casiraghi et al., Modeling of TDP-43 proteinopathy by chronic oxidative stress identifies rapamycin as beneficial in ALS patient-derived 2D and 3D iPSC models. Exp. Neurol. 383, 115057 (2025). 10.1016/j.expneurol.2024.115057.

14. Y. C. Liu, P. M. Chiang, K. J. Tsai, Disease animal models of TDP-43 proteinopathy and their pre-clinical applications. Int. J. Mol. Sci. 14, 20079–20111 (2013). 10.3390/ijms141020079.

15. R. M. Marton, S. P. PaLJca, Organoid and assembloid technologies for investigating cellular crosstalk in human brain development and disease. Trends Cell Biol. 30, 133–143 (2020). 10.1016/j.tcb.2019.11.004.

16. O. L. Eichmüller, J. A. Knoblich, Human cerebral organoids—a new tool for clinical neurology research. Nat. Rev. Neurol. 18, 661–680 (2022). 10.1038/s41582-022-00723-9.

17. Y. Sun et al., Brain-wide neuronal circuit connectome of human glioblastoma. Nature 641, 222–231 (2025). 10.1038/s41586-025-08634-7.

18. W. M. Babinchak et al., The role of liquid-liquid phase separation in aggregation of the TDP-43 low-complexity domain. J. Biol. Chem. 294, 6306–6317 (2019). 10.1074/jbc.RA118.007222.

19. H. J. Chen et al., RRM adjacent TARDBP mutations disrupt RNA binding and enhance TDP-43 proteinopathy. Brain 142, 3753–3770 (2019). 10.1093/brain/awz313.

20. J. Gu et al., Hsp70 chaperones TDP-43 in dynamic, liquid-like phase and prevents it from amyloid aggregation. Cell Res. 31, 1024–1027 (2021). 10.1038/s41422-021-00526-5.

21. Z. R. Grese et al., Specific RNA interactions promote TDP-43 multivalent phase separation and maintain liquid properties. EMBO Rep. 22, e53632 (2021). 10.15252/embr.202153632.

22. C. B. Pantazis et al., A reference human induced pluripotent stem cell line for large-scale collaborative studies. Cell Stem Cell 29, 1685–1702.e22 (2022). 10.1016/j.stem.2022.11.004.

23. D. M. Ramos et al., Tackling neurodegenerative diseases with genomic engineering: A new stem cell initiative from the NIH. Neuron 109, 1080–1083 (2021). 10.1016/j.neuron.2021.03.022.

24. Y. Zhang et al., Rapid single-step induction of functional neurons from human pluripotent stem cells. Neuron 78, 785–798 (2013). 10.1016/j.neuron.2013.05.029.

25. Y. Inukai et al., Abnormal phosphorylation of Ser409/410 of TDP-43 in FTLD-U and ALS. FEBS Lett. 582, 2899–2904 (2008). 10.1016/j.febslet.2008.07.027.

26. A. C. Raulin et al., Impact of APOE on amyloid and tau accumulation in argyrophilic grain disease and Alzheimer’s disease. Acta Neuropathol. Commun. 12, 25 (2024). 10.1186/s40478-024-01731-0.

27. V. I. Ko et al., CK1δ/ε kinases regulate TDP-43 phosphorylation and are therapeutic targets for ALS-related TDP-43 hyperphosphorylation. Neurobiol. Dis. 196, 106516 (2024). 10.1016/j.nbd.2024.106516.

28. A. Murakami et al., Old age amyotrophic lateral sclerosis and limbic TDP-43 pathology. Brain Pathol. 32, e13100 (2022). 10.1111/bpa.13100.

29. A. Bhaduri et al., Cell stress in cortical organoids impairs molecular subtype specification. Nature 578, 142–148 (2020). 10.1038/s41586-020-1962-0.

30. S. Porta et al., Patient-derived frontotemporal lobar degeneration brain extracts induce formation and spreading of TDP-43 pathology in vivo. Nat. Commun. 9, 4220 (2018). 10.1038/s41467-018-06548-9.

31. S. J. Barmada et al., Cytoplasmic mislocalization of TDP-43 is toxic to neurons and enhanced by a mutation associated with familial amyotrophic lateral sclerosis. J. Neurosci. 30, 639–649 (2010). 10.1523/JNEUROSCI.4988-09.2010.

32. R. Hasan et al., Transcriptomic analysis of frontotemporal lobar degeneration with TDP-43 pathology reveals cellular alterations across multiple brain regions. Acta Neuropathol. 143, 383–401 (2022). 10.1007/s00401-021-02399-9.

33. O. Britanova et al., Satb2 is a postmitotic determinant for upper-layer neuron specification in the neocortex. Neuron 57, 378–392 (2008). 10.1016/j.neuron.2007.12.028.

34. T. R. Young et al., Ten-m2 is required for the generation of binocular visual circuits. J. Neurosci. 33, 12490–12509 (2013). 10.1523/JNEUROSCI.4708-12.2013.

35. X. Zhang et al., Teneurins assemble into presynaptic nanoclusters that promote synapse formation via postsynaptic non-teneurin ligands. Nat. Commun. 13, 2297 (2022). 10.1038/s41467-022-29751-1.

36. X. Wang et al., The C. elegans homolog of human panic-disorder risk gene TMEM132D orchestrates neuronal morphogenesis through the WAVE-regulatory complex. Mol. Brain 14, 54 (2021). 10.1186/s13041-021-00767-w.

37. S. S. Pineda et al., Single-cell dissection of the human motor and prefrontal cortices in ALS and FTLD. Cell 187, 1971–1989.e16 (2024). 10.1016/j.cell.2024.02.031.

38. S. Wang, S. Sun, Translation dysregulation in neurodegenerative diseases: A focus on ALS. Mol. Neurodegener. 18, 58 (2023). 10.1186/s13024-023-00642-3.

39. S. Jin et al., Inference and analysis of cell-cell communication using CellChat. Nat. Commun. 12, 1088 (2021). 10.1038/s41467-021-21246-9.

40. H. Takahashi et al., Postsynaptic TrkC and presynaptic PTPσ function as a bidirectional excitatory synaptic organizing complex. Neuron 69, 287–303 (2011). 10.1016/j.neuron.2010.12.024.

41. C. Faux et al., PTPσ binds and dephosphorylates neurotrophin receptors and can suppress NGF-dependent neurite outgrowth from sensory neurons. Biochim. Biophys. Acta 1773, 1689–1700 (2007). 10.1016/j.bbamcr.2007.06.008.

42. R. Fernandez-Calle et al., Role of RPTPβ/ζ in neuroinflammation and microglia-neuron communication. Sci. Rep. 10, 20259 (2020). 10.1038/s41598-020-76415-5.

43. E. L. Van Nostrand et al., Robust, cost-effective profiling of RNA binding protein targets with single-end enhanced crosslinking and immunoprecipitation (seCLIP). Methods Mol. Biol. 1648, 177–200 (2017). 10.1007/978-1-4939-7204-3_14.

44. X. R. Ma et al., TDP-43 represses cryptic exon inclusion in the FTD-ALS gene UNC13A. Nature 603, 124–130 (2022). 10.1038/s41586-022-04424-7.

45. P. R. Mehta et al., The era of cryptic exons: Implications for ALS-FTD. Mol. Neurodegener. 18, 16 (2023). 10.1186/s13024-023-00608-5.

46. M.W. Baughn et al., Mechanism of STMN2 cryptic splice-polyadenylation and its correction for TDP-43 proteinopathies. Science 379, 1140–1149 (2023). doi:10.1126/science.abq5622.

47. A. L. Brown et al., TDP-43 loss and ALS-risk SNPs drive mis-splicing and depletion of UNC13A. Nature 603, 131–137 (2022). 10.1038/s41586-022-04436-3.

48. S. Shen et al., rMATS: Robust and flexible detection of differential alternative splicing from replicate RNA-Seq data. Proc. Natl. Acad. Sci. U.S.A. 111, E5593–E5601 (2014). 10.1073/pnas.1419161111.

49. J. R. Klim et al., ALS-implicated protein TDP-43 sustains levels of STMN2, a mediator of motor neuron growth and repair. Nat. Neurosci. 22, 167–179 (2019). 10.1038/s41593-018-0300-4.

50. Z. Melamed et al., Premature polyadenylation-mediated loss of stathmin-2 is a hallmark of TDP-43-dependent neurodegeneration. Nat. Neurosci. 22, 180–190 (2019). 10.1038/s41593-018-0293-z.

51. S. Cheedipudi et al., A fine balance: Epigenetic control of cellular quiescence by the tumor suppressor PRDM2/RIZ at a bivalent domain in the cyclin A gene. Nucleic Acids Res. 43, 6236–6256 (2015). 10.1093/nar/gkv567.

52. C. L. Bennett et al., Senataxin mutations elicit motor neuron degeneration phenotypes and yield TDP-43 mislocalization in ALS4 mice and human patients. Acta Neuropathol. 136, 425–443 (2018). 10.1007/s00401-018-1852-9.

53. A. Menga et al., The mitochondrial aspartate/glutamate carrier isoform 1 gene expression is regulated by CREB in neuronal cells. Int. J. Biochem. Cell Biol. 60, 157–166 (2015). 10.1016/j.biocel.2015.01.004.

54. F. Palmieri, The mitochondrial transporter family SLC25: Identification, properties and physiopathology. Mol. Aspects Med. 34, 465–484 (2013). 10.1016/j.mam.2012.05.005.

55. A. Devarajan et al., Paraoxonase 2 deficiency alters mitochondrial function and exacerbates the development of atherosclerosis. Antioxid. Redox Signal. 14, 341–351 (2011). 10.1089/ars.2010.3430.

56. M. D. Brand, T. C. Esteves, Physiological functions of the mitochondrial uncoupling proteins UCP2 and UCP3. Cell Metab. 2, 85–93 (2005). 10.1016/j.cmet.2005.06.002.

57. J. St-Pierre et al., Suppression of reactive oxygen species and neurodegeneration by the PGC-1 transcriptional coactivators. Cell 127, 397–408 (2006). 10.1016/j.cell.2006.09.024.

58. Y. Gonda, T. Namba, C. Hanashima, Beyond axon guidance: Roles of Slit-Robo signaling in neocortical formation. Front. Cell Dev. Biol. 8, 607415 (2020). 10.3389/fcell.2020.607415.

59. N. Neelagandan et al., TDP-43 enhances translation of specific mRNAs linked to neurodegenerative disease. Nucleic Acids Res. 47, 341–361 (2019). 10.1093/nar/gky972.

60. R. Barchiesi et al., An epigenetic mechanism for over-consolidation of fear memories. Mol. Psychiatry 27, 4893–4904 (2022). 10.1038/s41380-022-01758-6.

61. E. Barbier et al., Dependence-induced increase of alcohol self-administration and compulsive drinking mediated by the histone methyltransferase PRDM2. Mol. Psychiatry 22, 1746–1758 (2017). 10.1038/mp.2016.131.

62. I. Niebroj-Dobosz et al., Matrix metalloproteinases and their tissue inhibitors in serum and cerebrospinal fluid of patients with amyotrophic lateral sclerosis. Eur. J. Neurol. 17, 226–231 (2010). 10.1111/j.1468-1331.2009.02775.x.

63. J. Lin et al., Key molecules and pathways underlying sporadic amyotrophic lateral sclerosis: Integrated analysis on gene expression profiles of motor neurons. Front. Genet. 11, 578143 (2020). 10.3389/fgene.2020.578143.

64. S. Apolloni et al., Extracellular matrix remodeling in motor neuron diseases. Int. J. Mol. Sci. 26, 11376 (2025). 10.3390/ijms262311376.

65. T. Cunha-Oliveira et al., Oxidative stress in amyotrophic lateral sclerosis: Pathophysiology and opportunities for pharmacological intervention. Oxid. Med. Cell. Longev. 2020, 5021694 (2020). 10.1155/2020/5021694.

66. M. S. Fernandopulle et al., Transcription factor–mediated differentiation of human iPSCs into neurons. Curr. Protoc. Cell Biol. 79, e51 (2018). 10.1002/cpcb.51.

67. H. N. Nguyen, Generation of iPSC-derived brain organoids for drug testing and toxicological evaluation. Methods Mol. Biol. 2474, 93–105 (2022). 10.1007/978-1-0716-2213-1_10.

68. A. S. Buchman, T. Wang, L. Yu, S. E. Leurgans, J. A. Schneider, D. A. Bennett, Brain pathologies are associated with both the rate and variability of declining motor function in older adults. Acta Neuropathol. 140, 587–589 (2020). 10.1007/s00401-020-02212-z.

69. T. Wu et al., clusterProfiler 4.0: A universal enrichment tool for interpreting omics data. Innovation (Camb.) 2, 100141 (2021). 10.1016/j.xinn.2021.100141.

70. T. Smith, A. Heger, I. Sudbery, UMI-tools: Modeling sequencing errors in unique molecular identifiers to improve quantification accuracy. Genome Res. 27, 491–499 (2017). 10.1101/gr.209601.116.

71. R. Hubley et al., The Dfam database of repetitive DNA families. Nucleic Acids Res. 44, D81–D89 (2016). 10.1093/nar/gkv1272.

72. A. Dobin et al., STAR: Ultrafast universal RNA-seq aligner. Bioinformatics 29, 15–21 (2013). 10.1093/bioinformatics/bts635.

73. M. T. Lovci et al., Rbfox proteins regulate alternative mRNA splicing through evolutionarily conserved RNA bridges. Nat. Struct. Mol. Biol. 20, 1434–1442 (2013). 10.1038/nsmb.2699.

74. A. Frankish et al., GENCODE: Reference annotation for the human and mouse genomes in 2023. Nucleic Acids Res. 51, D942–D949 (2023). 10.1093/nar/gkac1071.

75. S. Heinz et al., Simple combinations of lineage-determining transcription factors prime cis-regulatory elements required for macrophage and B cell identities. Mol. Cell 38, 576–589 (2010). 10.1016/j.molcel.2010.05.004.

76. G. H. Putri, S. Anders, P. T. Pyl, J. E. Pimanda, F. Zanini, Analysing high-throughput sequencing data in Python with HTSeq 2.0. Bioinformatics 38, 2943–2945 (2022). 10.1093/bioinformatics/btac166.

77. M. I. Love, W. Huber, S. Anders, Moderated estimation of fold change and dispersion for RNA-seq data with DESeq2. Genome Biol. 15, 550 (2014). 10.1186/s13059-014-0550-8.

