## supplemental materials for "An ALS-associated TARDBP mutation drives cryptic exon inclusion and RNA dysregulation"

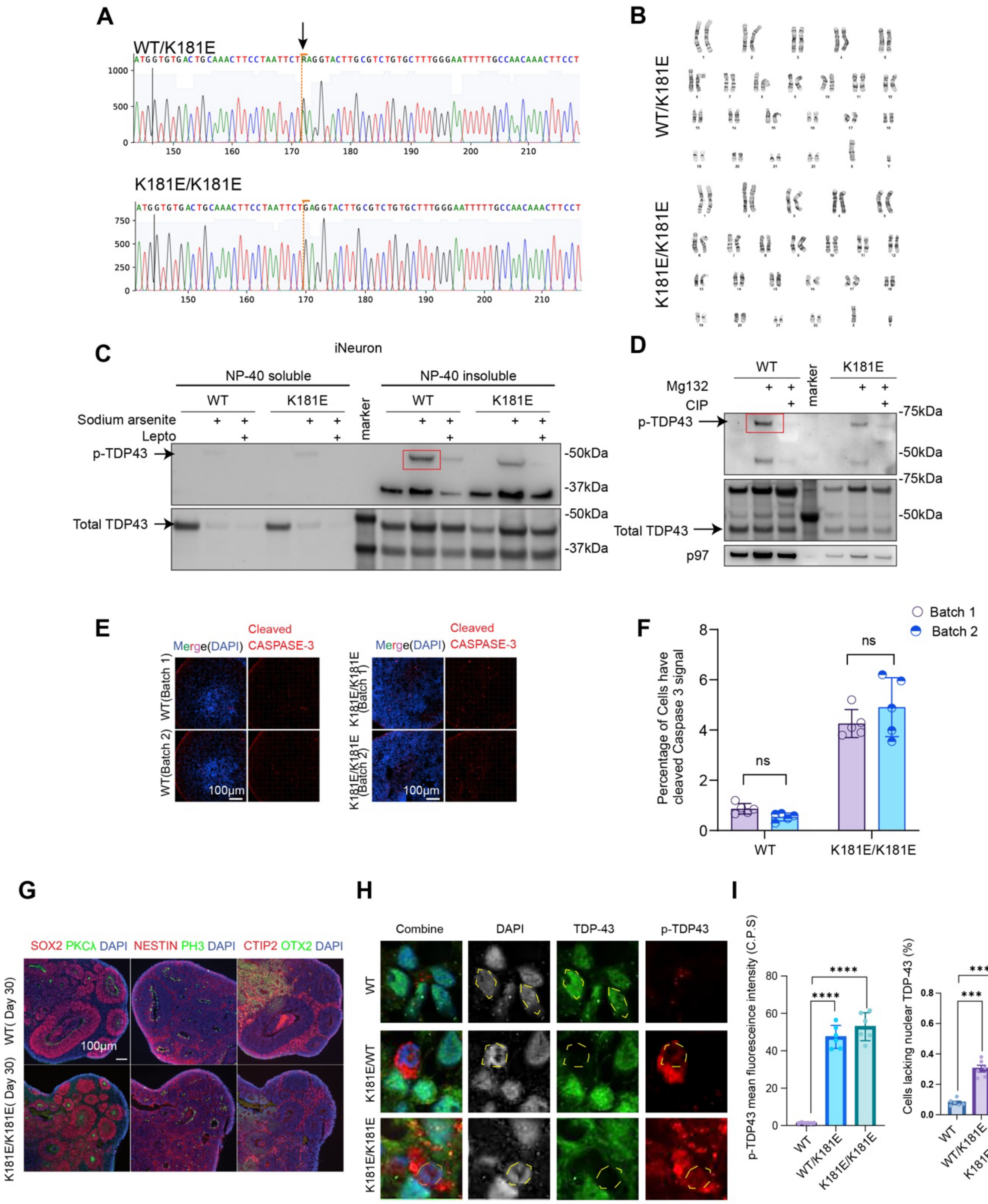

A
WT/K181E
K181E/K181E
B
WT/K181E
K181E/K181E
C
iNeuron
NP-40 soluble
NP-40 insoluble
WT
K181E
Sodium arsenite
Lepto
marker
p-TDP43
Total TDP43
-50kDa
-37kDa
D
WT
K181E
Mg132
CIP
marker
p-TDP43
Total TDP43
p97
-75kDa
-50kDa
E
Merge(DAPI)
Cleaved CASPASE-3
WT(Batch 1)
WT(Batch 2)
K181E/K181E (Batch 1)
K181E/K181E (Batch 2)
100μm
F
Batch 1
Batch 2
Percentage of Cells have cleaved Caspase 3 signal
ns
WT
K181E/K181E
G
SOX2 PKCλ DAPI
NESTIN PH3 DAPI
CTIP2 OTX2 DAPI
WT( Day 30)
K181E/K181E( Day 30)
100μm
H
Combine
DAPI
TDP-43
p-TDP43
WT
K181E/WT
K181E/K181E
I
p-TDP43 mean fluorescence intensity (C.P.S)
Cells lacking nuclear TDP-43 (%)
WT
WT/K181E
K181E/K181E
****
***

**Figure S1 Validation of TDP-43 K181E mutation knock-in and characterization of TDP-43 hyperphosphorylation in iNeuron derived excitatory neuron and forebrain organoid**

**(A)** Sanger sequencing confirms iPSC clones with one allele of TDP-43 K181E (top) or homozygous for TDP-43 K181E (bottom).
**(B)** Chromosome analysis confirms the correct karyotype for TDP-43 K181E clones.
(**C**) Hyperphosphorylation of endogenous TDP-43 (arrow and red box) was only detected in NP-40 insoluble fraction of mature $i^3$Neurons when oxidative stress was induced by sodium arsenite and was suppressed when cells were co-treated with Leptomycin B (Lepto). $i^3$Neurons of the indicated genotypes were treated with DMSO as a control, or with sodium arsenite (10 μM) or sodium arsenite and Leptomycin B (200 nM) at day 15 in differentiation. Cells were harvested at day 17 and lysed in a buffer containing NP40. NP40 insoluble fractions were further treated with an SDS-containing buffer before immunoblotting analysis.
(**D**) The specificity of the phospho-TDP-43 antibody was tested in HEK293T cells transfected with plasmids expressing ΔNLS-TDP-43-mGreenLantern and ΔNLS-TDP-43 (K181E)-mGreenLantern. Where indicated, cells were treated with the proteasome inhibitor MG132 (10 μM). A fraction of the lysate was treated with a phosphatase (CIP) before immunoblotting.
**(E)** Immunostaining of cleaved Caspase-3 in two individual batches of WT organoids (left) and TDP-43 K181E HM mutant organoids (right). n= 3-5 organoids, 2-3 iPSC clones, two individual batches.
**(F)** Quantification of percentage of cells have cleaved Caspase-3 signal in e, n= 3-5 organoids, 2-3 iPSC clones, two individual batches. Two-way ANOVA. Error bars indicate mean ± s.e.m, ns (no significant).
**(G)** Immunostaining of wild-type (WT) and K181E HM forebrain organoids at day 30 showed expression patterns consistent with early forebrain development. PKCλ labeled the apical surface of neuroepithelial structures, indicating ventricular zone-like organization. NESTIN marked neural stem and progenitor cells. PH3 labeled dividing neural stem and progenitor cells in mitosis. CTIP2 marked deep-layer cortical projection neurons, mostly those in layer V. OTX2 indicated anterior forebrain identity. SOX2 labeled neural stem and progenitor cells.

**(H)** Immunostaining of WT, TDP-43 WT/K181E, and TDP-43 K181E/K181E organoids with antibodies recognizing phospho-TDP-43 (Ser 409/410) and total TDP-43. Nucleus is highlighted by dashed yellow line. Cells with strong p-TDP-43 signal showed loss of nuclear total TDP-43 signal, whereas cells with nuclear total TDP-43 staining do not have p-TDP43 signal, supporting nuclear TDP-43 depletion in cells with TDP-43 hyperphosphorylation. DAPI was used to label nuclei.

**(I)** Quantification of phospho-TDP-43 mean fluorescence intensity and the percentage of cells lacking nuclear TDP-43 in WT, TDP-43 WT/K181E, and TDP-43 K181E/K181E organoids shown in (H). One-way ANOVA. Error bars indicate mean ± s.e.m.; ***$P < 0.001$, ****$P < 0.0001$.

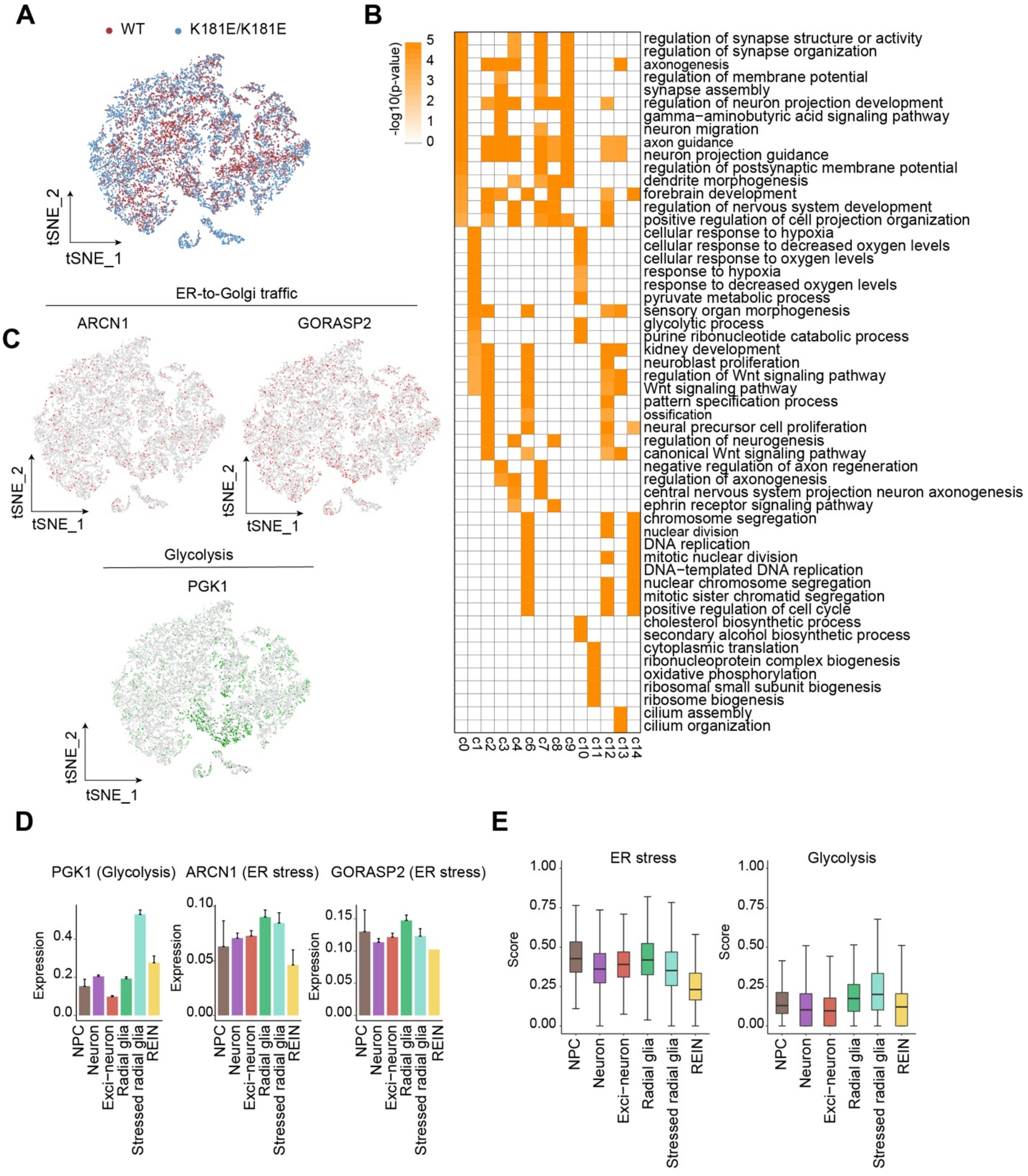
A
WT
K181E/K181E
tSNE_2
tSNE_1
B
-log10(p-value)
5
4
3
2
1
0
regulation of synapse structure or activity
regulation of synapse organization
axonogenesis
regulation of membrane potential
synapse assembly
regulation of neuron projection development
gamma−aminobutyric acid signaling pathway
neuron migration
axon guidance
neuron projection guidance
regulation of postsynaptic membrane potential
dendrite morphogenesis
forebrain development
regulation of nervous system development
positive regulation of cell projection organization
cellular response to hypoxia
cellular response to decreased oxygen levels
cellular response to oxygen levels
response to hypoxia
response to decreased oxygen levels
pyruvate metabolic process
sensory organ morphogenesis
glycolytic process
purine ribonucleotide catabolic process
kidney development
neuroblast proliferation
regulation of Wnt signaling pathway
Wnt signaling pathway
pattern specification process
ossification
neural precursor cell proliferation
regulation of neurogenesis
canonical Wnt signaling pathway
negative regulation of axon regeneration
regulation of axonogenesis
central nervous system projection neuron axonogenesis
ephrin receptor signaling pathway
chromosome segregation
nuclear division
DNA replication
mitotic nuclear division
DNA−templated DNA replication
nuclear chromosome segregation
mitotic sister chromatid segregation
positive regulation of cell cycle
cholesterol biosynthetic process
secondary alcohol biosynthetic process
cytoplasmic translation
ribonucleoprotein complex biogenesis
oxidative phosphorylation
ribosomal small subunit biogenesis
ribosome biogenesis
cilium assembly
cilium organization
c0 c1 c2 c3 c4 c6 c7 c8 c9 c10 c11 c12 c13 c14
ER-to-Golgi traffic
C
ARCN1
GORASP2
tSNE_2
tSNE_1
Glycolysis
PGK1
D
PGK1 (Glycolysis)
ARCN1 (ER stress)
GORASP2 (ER stress)
Expression
0.0
0.2
0.4
0.00
0.05
0.10
0.15
NPC
Neuron
Exci−neuron
Radial glia
Stressed radial glia
REIN
E
ER stress
Glycolysis
Score
0.00
0.25
0.50
0.75
1.00

**Figure S2 Single-cell transcriptomic analysis reveals genotype-specific clustering and differential enrichment of ER stress and glycolytic pathways.**

**(A)** T-distributed Stochastic Neighbor Embedding (t-SNE) of 16,549 single cells colored by sample genotypes.

**(B)** Heatmap showing the GO enriched pathways in each cell cluster. The significance was determined by adjusted p-value <0.05. Top 5 biological pathways for each cell cluster were displayed based on p-value.

**(C)** Feature plots showing the expression of genes involved in ER-to-Golgi traffic (ARCN1 and GORASP2) and glycolysis (PGK1).

**(D)** Bar graphs showing the relative expression of the indicated genes in each cell type. Shown are mean ± s.e.m.

**(E)** Box plots showing pathway score of ER stress (left) and glycolysis (right). Genes involved in the pathways were obtained from the GSEA database. The pathway scores were calculated as average expression of the genes.

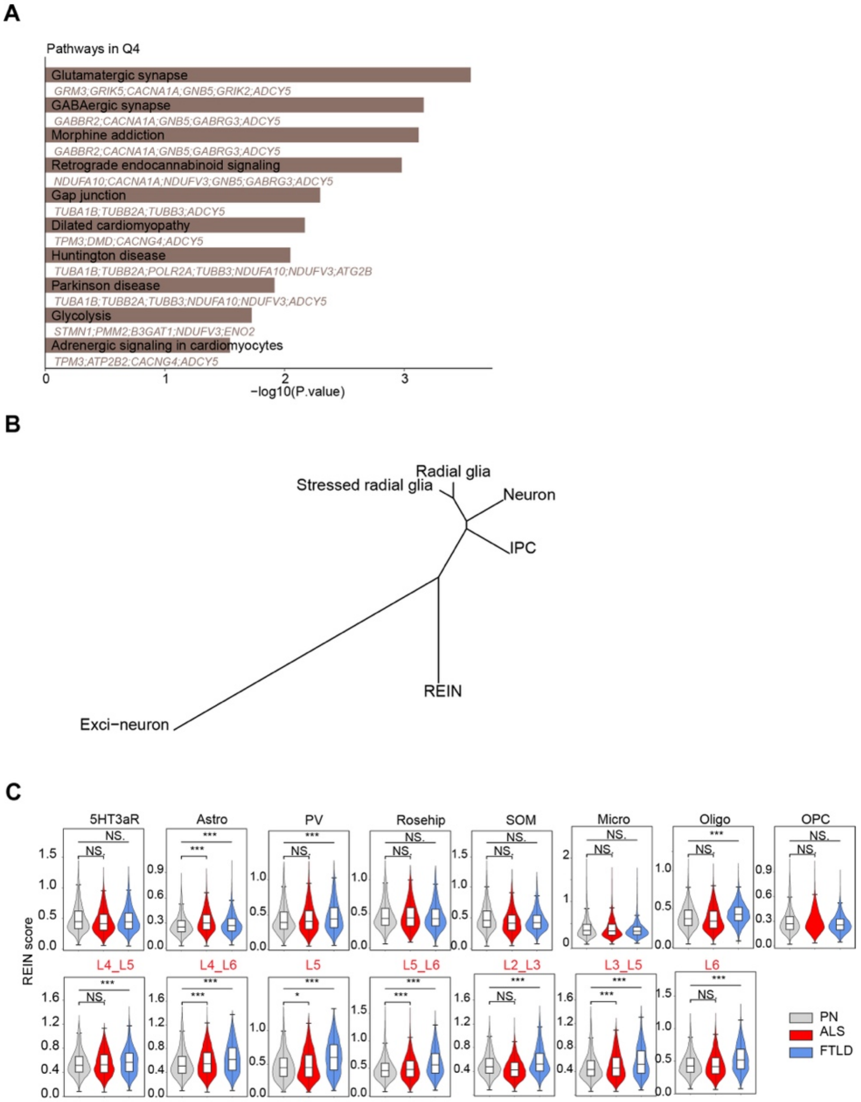


**Figure S3 KEGG pathway enrichment for genes in Q4 and cell type specific REIN scores in ALS/FTD patients and hierarchical clustering of organoid cell types.**

**(A)** Top enriched KEGG pathways of the genes in Q4 (n=144) from (Fig. 2A).

**(B)** Hierarchical clustering of annotated cell types from organoid scRNA-seq data using Pearson distance calculated from cell type-averaged transcriptional profiles. REINs clustered in relation to neuronal populations and were separated from glial and progenitor-enriched populations, supporting interpretation of REINs as a neuronal transcriptional state.

(**C**) Violin plots of REIN scores in different cell types from ALS/FTD patients (* $p < 0.05$, *** $p < 0.001$ by two-tailed unpaired Student's t-test. NS, non-significant). The REIN score is calculated as the average expression of REIN-related genes (Supplementary Table S2). (L4-L6: Cortical layer L4-L6, Micro: microglia, Oligo: oligodendrocyte, OPC: oligodendrocyte progenitor cells, SOM: **somatostatin, Rosehip: rosehip neuron, PV: PV neuron, Astro: astrocyte, 5HT3aR: 5HT3aR GABAergic interneuron**).

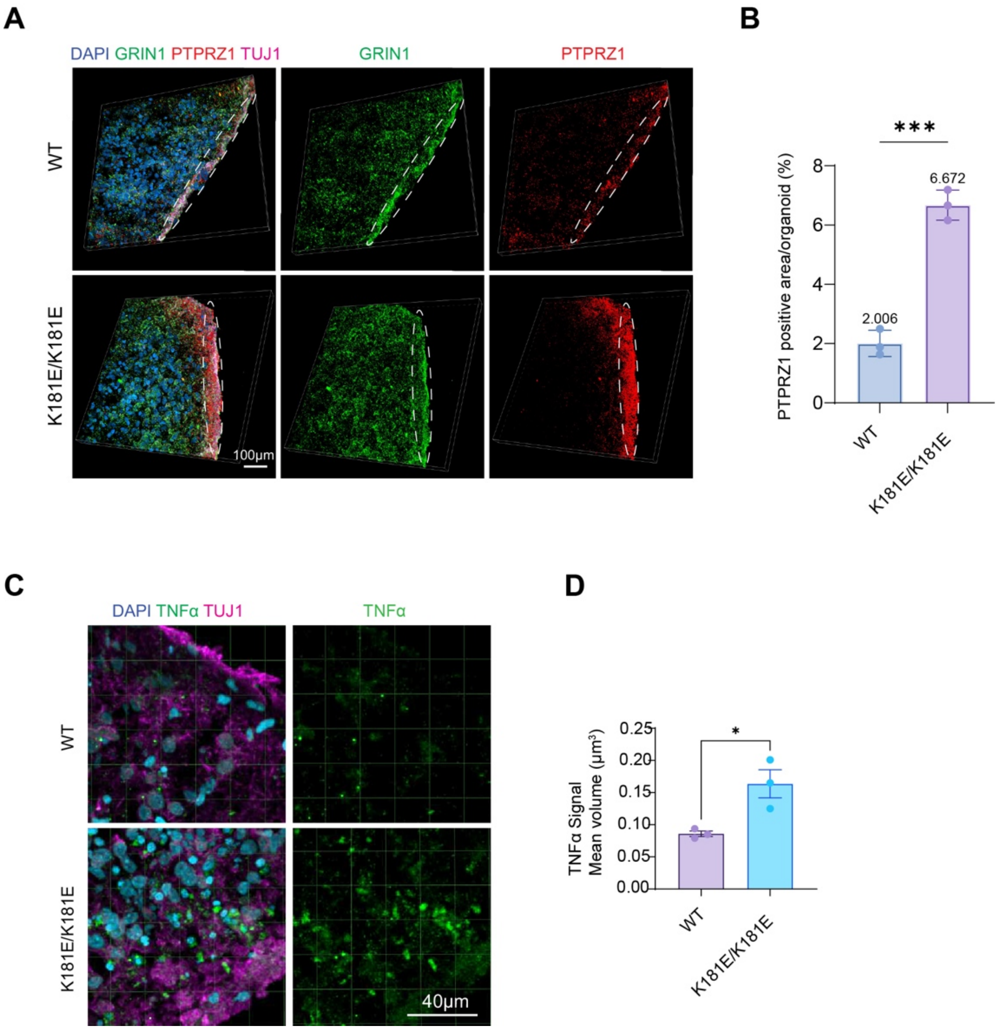


**Figure S4 Increased PTPRZ1 and TNF-α expression in TDP-43 K181E Homo mutant organoid.**

**(A)** Representative confocal fluorescence images of immuno-stained organoid sections showing upregulated expression of PTPRZ1(red) in organoids bearing two alleles of K181E (K181E/K181E). Organoids were also stained for neuronal markers TUJ1 (magenta) and GRIN1 (green) as controls. Scale bar, 100 μm. n= 3-5 organoids, 2-3 iPSC clones, two individual batches.

**(B)** Quantification of PTPRZ1 positive area per organoid in 3D fluorescence images of organoids with the indicated genotypes. Error bars indicate mean ± s.e.m., **** $p < 0.0001$ by unpaired t-test, n=3 organoids.

**(C)** Representative confocal fluorescence images of immuno-stained organoid sections showing upregulated expression of TNF-α (green) in organoids bearing two alleles of K181E

(K181E/K181E). Organoids were also stained for neuronal marker TUJ1 (magenta) Scale bar, 100 µm. n= 3-5 organoids, 2-3 iPSC clones, two individual batches.

**(D)** Quantification of TNF-α positive mean spot volume in 3D fluorescence images of organoids with the indicated genotypes. Error bars indicate mean ± s.e.m., **** $p < 0.0001$ by unpaired t-test, n=3 organoids.

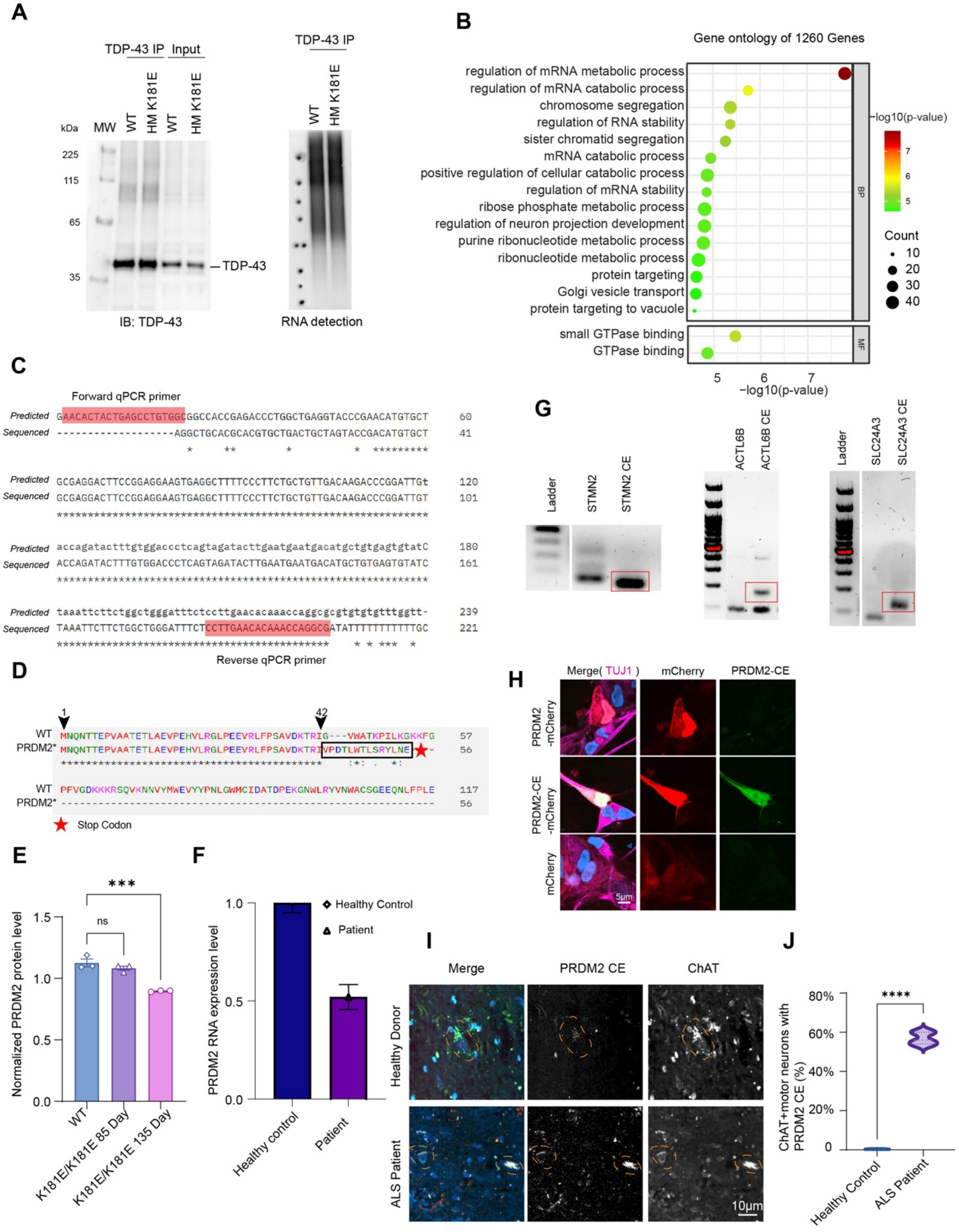

A
TDP-43 IP
Input
kDa
MW
WT
HM K181E
WT
HM K181E
225
115
65
35
TDP-43
IB: TDP-43
TDP-43 IP
WT
HM K181E
RNA detection
B
Gene ontology of 1260 Genes
regulation of mRNA metabolic process
regulation of mRNA catabolic process
chromosome segregation
regulation of RNA stability
sister chromatid segregation
mRNA catabolic process
positive regulation of cellular catabolic process
regulation of mRNA stability
ribose phosphate metabolic process
regulation of neuron projection development
purine ribonucleotide metabolic process
ribonucleotide metabolic process
protein targeting
Golgi vesicle transport
protein targeting to vacuole
small GTPase binding
GTPase binding
BP
MF
−log10(p-value)
7
6
5
Count
10
20
30
40
5
6
7
−log10(p-value)
C
Forward qPCR primer
Predicted GAACACTACTGAGCCTGTGGCGGCCACCGAGACCCTGGCTGAGGTACCCGAACATGTGCT 60
Sequenced ------------------AGGCTGCACGCACGTGCTGACTGCTAGTACCGACATGTGCT 41
Predicted GCGAGGACTTCCGGAGGAAGTGAGGCTTTTCCCTTCTGCTGTTGACAAGACCCGGATTGt 120
Sequenced GCGAGGACTTCCGGAGGAAGTGAGGCTTTTCCCTTCTGCTGTTGACAAGACCCGGATTGT 101
Predicted accagatactttgtggaccctcagtagatacttgaatgaatgacatgctgtgagtgtatC 180
Sequenced ACCAGATACTTTGTGGACCCTCAGTAGATACTTGAATGAATGACATGCTGTGAGTGTATC 161
Predicted taaattcttctggctgggatttctccttgaacacaaaccaggcgcgtgtgtgtttggtt- 239
Sequenced TAAATTCTTCTGGCTGGGATTTCTCCTTGAACACAAACCAGGCGATATTTTTTTTTTTGC 221
Reverse qPCR primer
D
1
42
WT MNQNTTEPVAATETLAEVPEHVLRGLPEEVRLFPSAVDKTRIG---VWATKPILKGKKFG 57
PRDM2* MNQNTTEPVAATETLAEVPEHVLRGLPEEVRLFPSAVDKTRIVPDTLWTLSRYLNE- 56
WT PFVGDKKKRSQVKNNVYMWEVYYPNLGWMCIDATDPEKGNWLRYVNWACSGEEQNLFPLE 117
PRDM2* ------------------------------------------------------------ 56
Stop Codon
E
***
ns
Normalized PRDM2 protein level
1.5
1.0
0.5
0.0
WT
K181E/K181E 85 Day
K181E/K181E 135 Day
F
Healthy Control
Patient
PRDM2 RNA expression level
1.0
0.5
0.0
Healthy control
Patient
G
Ladder
STMN2
STMN2 CE
ACTL6B
ACTL6B CE
Ladder
SLC24A3
SLC24A3 CE
H
Merge( TUJ1 )
mCherry
PRDM2-CE
PRDM2 -mCherry
PRDM2-CE -mCherry
mCherry
5µm
I
Merge
PRDM2 CE
ChAT
Healthy Donor
ALS Patient
10µm
J
****
ChAT+motor neurons with PRDM2 CE (%)
80%
60%
40%
20%
0
Healthy Control
ALS Patient

**Figure S5 TDP-43 RNA immunoprecipitation, Gene Ontology analysis of WT-specific bound mRNAs, and validation of PRDM2 cryptic exon inclusion with reduced protein levels in mutant organoids.**

**(A)** Immunoblotting analysis of immunoprecipitated (IP) samples: 10% of IP samples and 1% of input samples were fractionated on a NuPAGE 4-12% Bis-Tris protein gel; Proteins were blotted by anti-TDP43 antibodies (left); RNA in immunoprecipitated samples were visualized by chemiluminescent nucleic acid detection kit (right).
**(B)** Gene Ontology analysis of the mRNAs that only bind WT TDP-43 (Fig. 4b, n=1260) by the eCLIP analysis.
**(C)** DNA sequence alignment reveals the detected CE sequence by Sanger sequencing. Sanger sequencing results of CE fragment amplified by PCR using cDNA as the template. Reference: Predicted CE containing sequence by rMATs. Primers used in qPCR were highlighted in red. Identified CE is indicated by small letters.
**(D)** PRDM2 protein sequence alignment between WT and the translation product of mis-spliced PRDM2 variant (PRDM2*).
**(E)** Normalized PRDM2 protein level in organoids with the indicated genotypes as determined by mass spectrometry. Error bars indicate mean ± s.e.m., one-way ANOVA; **** $p < 0.0001$, n=3 organoids.
**(F)** Normalized PRDM2 RNA level in healthy control and FTD patients, as determined by bulk RNAseq. Error bars indicate mean ± s.e.m., unpaired t-test; **** $p < 0.0001$.
**(G)** RT-PCR validation of selected cryptic exon events identified by RNA-seq/rMATS analysis. Primer pairs were designed to specifically amplify cryptic exon-containing transcripts, with the reverse primer located within the predicted cryptic exon sequence. CE-containing PCR products were confirmed for STMN2 (left), ACTL6B(middle) and SLC24A3 (right) in K181E/K181E organoids compared with WT control. Red boxes indicate PCR products corresponding to the predicted CE-containing amplicons. Additional bands outside the expected CE amplicon size were not used for interpretation and likely reflect nonspecific RT-PCR products or low-abundance precursor RNA.
**(H)** Validation of PRDM2 cryptic exon peptide antibody specificity in iPSC-derived motor neurons. Cells were infected for 48 h with lentiviruses expressing mCherry-tagged WT PRDM2, mCherry-tagged PRDM2 containing the cryptic exon-encoded sequence, or mCherry alone under the control of the neuronal synapsin promoter. Cells were stained with antibodies against TUJ1 and the PRDM2 CE peptide. PRDM2 CE-peptide immunoreactivity was detected in

PRDM2-CE-mCherry–expressing neurons, but not in neurons expressing WT PRDM2-mCherry or mCherry alone, supporting specificity of the antibody for the CE-encoded neoepitope. Scale bar, 5 μm.
**(I)** Representative immunofluorescence images of postmortem spinal cord sections from two independent healthy controls and three ALS patients stained with antibodies against choline acetyltransferase (ChAT) to label spinal motor neurons and PRDM2 cryptic exon-peptide (PRDM2 CE-peptide). A total of 8 sections were analyzed. Nuclei are stained with DAPI. In healthy control samples, PRDM2 CE-peptide signal is minimal or absent in ChAT-positive motor neurons, whereas ALS patient samples show detectable PRDM2 CE-peptide signal in ChAT-positive cells. Scale bars are indicated.

**(J)** Quantification (**I**) of the percentage of ChAT-positive motor neurons that are positive for PRDM2 cryptic exon-peptide signal in healthy donors and ALS patients. Each data point represents an individual spinal cord section (sections nested within donors). Error bars indicate mean ± s.e.m., **** $p < 0.0001$ unpaired Student's t-test, N=8.

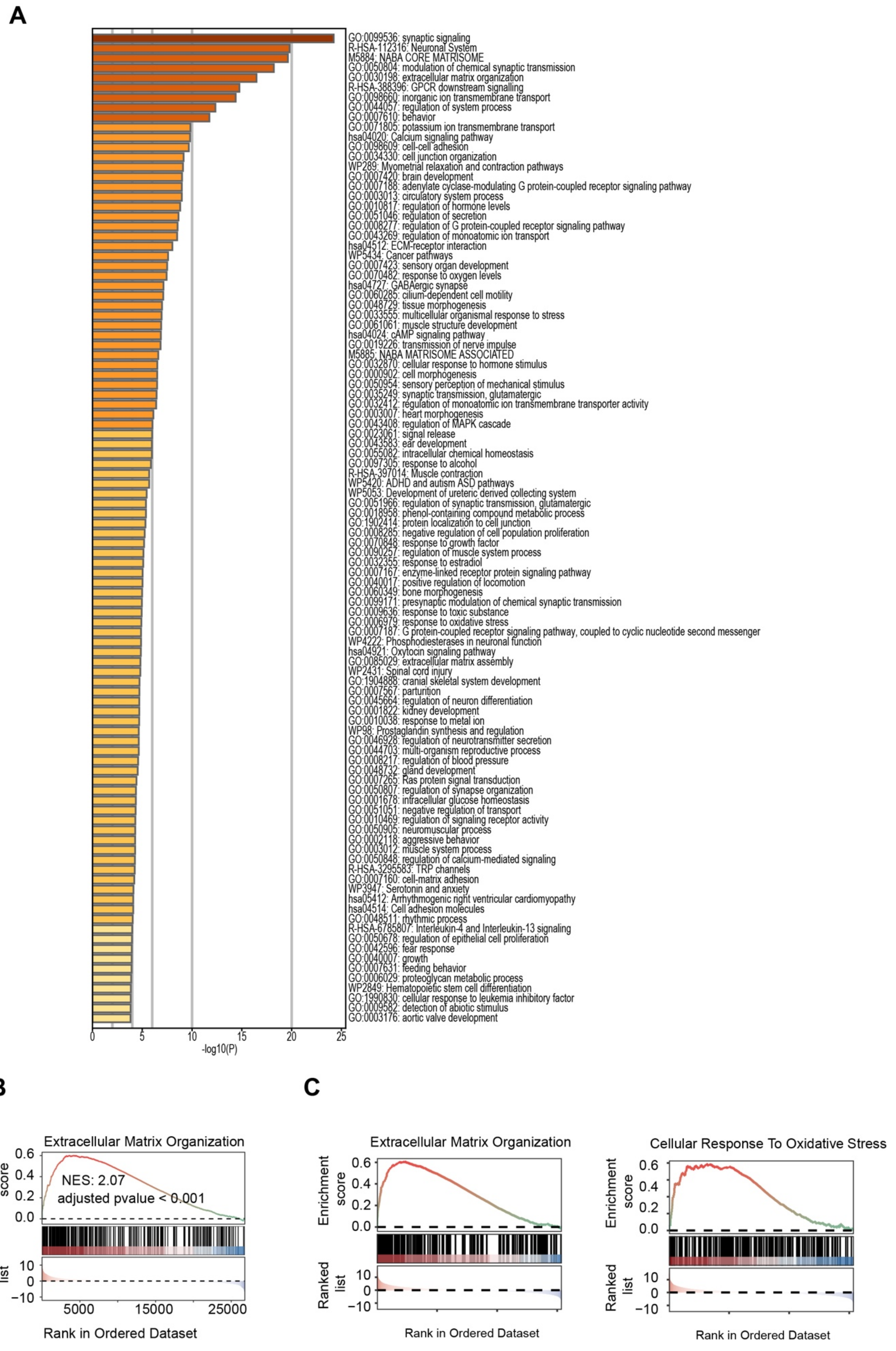

A
GO:0099536: synaptic signaling
R-HSA-112316: Neuronal System
M5884: NABA CORE MATRISOME
GO:0050804: modulation of chemical synaptic transmission
GO:0030198: extracellular matrix organization
R-HSA-388396: GPCR downstream signalling
GO:0098660: inorganic ion transmembrane transport
GO:0044057: regulation of system process
GO:0007610: behavior
GO:0071805: potassium ion transmembrane transport
hsa04020: Calcium signaling pathway
GO:0098609: cell-cell adhesion
GO:0034330: cell junction organization
WP289: Myometrial relaxation and contraction pathways
GO:0007420: brain development
GO:0007188: adenylate cyclase-modulating G protein-coupled receptor signaling pathway
GO:0003013: circulatory system process
GO:0010817: regulation of hormone levels
GO:0051046: regulation of secretion
GO:0008277: regulation of G protein-coupled receptor signaling pathway
GO:0043269: regulation of monoatomic ion transport
hsa04512: ECM-receptor interaction
WP5434: Cancer pathways
GO:0007423: sensory organ development
GO:0070482: response to oxygen levels
hsa04727: GABAergic synapse
GO:0060285: cilium-dependent cell motility
GO:0048729: tissue morphogenesis
GO:0033555: multicellular organismal response to stress
GO:0061061: muscle structure development
hsa04024: cAMP signaling pathway
GO:0019226: transmission of nerve impulse
M5885: NABA MATRISOME ASSOCIATED
GO:0032870: cellular response to hormone stimulus
GO:0000902: cell morphogenesis
GO:0050954: sensory perception of mechanical stimulus
GO:0035249: synaptic transmission, glutamatergic
GO:0032412: regulation of monoatomic ion transmembrane transporter activity
GO:0003007: heart morphogenesis
GO:0043408: regulation of MAPK cascade
GO:0023061: signal release
GO:0043583: ear development
GO:0055082: intracellular chemical homeostasis
GO:0097305: response to alcohol
R-HSA-397014: Muscle contraction
WP5420: ADHD and autism ASD pathways
WP5053: Development of ureteric derived collecting system
GO:0051966: regulation of synaptic transmission, glutamatergic
GO:0018958: phenol-containing compound metabolic process
GO:1902414: protein localization to cell junction
GO:0008285: negative regulation of cell population proliferation
GO:0070848: response to growth factor
GO:0090257: regulation of muscle system process
GO:0032355: response to estradiol
GO:0007167: enzyme-linked receptor protein signaling pathway
GO:0040017: positive regulation of locomotion
GO:0060349: bone morphogenesis
GO:0099171: presynaptic modulation of chemical synaptic transmission
GO:0009636: response to toxic substance
GO:0006979: response to oxidative stress
GO:0007187: G protein-coupled receptor signaling pathway, coupled to cyclic nucleotide second messenger
WP4222: Phosphodiesterases in neuronal function
hsa04921: Oxytocin signaling pathway
GO:0085029: extracellular matrix assembly
WP2431: Spinal cord injury
GO:1904888: cranial skeletal system development
GO:0007567: parturition
GO:0045664: regulation of neuron differentiation
GO:0001822: kidney development
GO:0010038: response to metal ion
WP98: Prostaglandin synthesis and regulation
GO:0046928: regulation of neurotransmitter secretion
GO:0044703: multi-organism reproductive process
GO:0008217: regulation of blood pressure
GO:0048732: gland development
GO:0007265: Ras protein signal transduction
GO:0050807: regulation of synapse organization
GO:0001678: intracellular glucose homeostasis
GO:0051051: negative regulation of transport
GO:0010469: regulation of signaling receptor activity
GO:0050905: neuromuscular process
GO:0002118: aggressive behavior
GO:0003012: muscle system process
GO:0050848: regulation of calcium-mediated signaling
R-HSA-3295583: TRP channels
GO:0007160: cell-matrix adhesion
WP3947: Serotonin and anxiety
hsa05412: Arrhythmogenic right ventricular cardiomyopathy
hsa04514: Cell adhesion molecules
GO:0048511: rhythmic process
R-HSA-6785807: Interleukin-4 and Interleukin-13 signaling
GO:0050678: regulation of epithelial cell proliferation
GO:0042596: fear response
GO:0040007: growth
GO:0007631: feeding behavior
GO:0006029: proteoglycan metabolic process
WP2849: Hematopoietic stem cell differentiation
GO:1990830: cellular response to leukemia inhibitory factor
GO:0009582: detection of abiotic stimulus
GO:0003176: aortic valve development
0
5
10
15
20
25
-log10(P)
B
Extracellular Matrix Organization
Enrichment score
0.6
0.4
0.2
0.0
NES: 2.07
adjusted pvalue < 0.001
Ranked list
10
0
-10
5000
15000
25000
Rank in Ordered Dataset
C
Extracellular Matrix Organization
Enrichment score
0.6
0.4
0.2
0.0
Ranked list
10
0
-10
Rank in Ordered Dataset
Cellular Response To Oxidative Stress
Enrichment score
0.6
0.4
0.2
0.0
Ranked list
10
0
-10
Rank in Ordered Dataset

**Figure S6 Shared transcriptional enrichment of oxidative stress and extracellular matrix pathways across PRDM2 knockdown motor neurons, TDP-43 K181E organoids, and patient datasets**

**(A)** Gene Ontology and pathway enrichment analysis of genes upregulated in TDP-43 K181E forebrain organoid. Bars represent significantly enriched biological processes and signaling pathways ranked by −log10(*P* value). Enriched categories include cellular responses to oxidative stress, extracellular matrix organization, synaptic signaling, and cell adhesion–related pathways.

**(B)** Gene set enrichment analysis (GSEA) showing significant enrichment of the Extracellular Matrix Organization gene set in PRDM2 knockdown motor neurons (normalized enrichment score (NES) = 2.07, adjusted $P < 0.001$).

**(C)** GSEA showing enrichment of Extracellular Matrix Organization and Cellular Response to Oxidative Stress gene sets in transcriptomic profiles derived from FTD patient datasets.

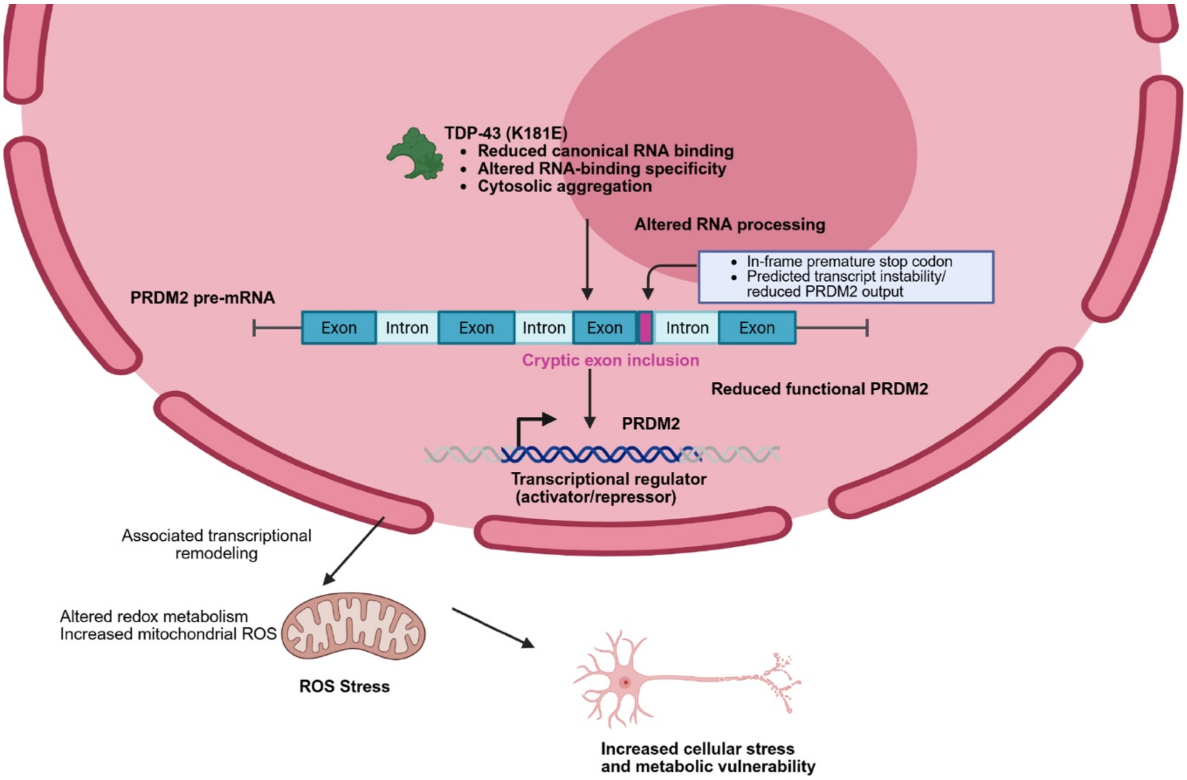


**Figure S7. Conceptual model linking TDP-43 K181E–associated RNA mis-splicing of PRDM2 to convergent neuronal stress responses**

Schematic model illustrating how the ALS/FTD-associated TDP-43 K181E mutation alters RNA binding and processing, leading to cryptic exon inclusion in PRDM2 pre-mRNA. Inclusion of the cryptic exon introduces an in-frame premature stop codon, resulting in predicted transcript instability and reduced functional PRDM2 output. Reduced PRDM2, a transcriptional regulator with both activating and repressive functions, is associated with activation of mitochondrial oxidative stress pathway. These convergent molecular changes are proposed to increase cellular stress and metabolic vulnerability in neurons, providing a mechanistic framework linking TDP-43–dependent RNA mis-splicing to downstream stress responses relevant to ALS and FTD pathology.

**Supplementary tables:**

Table S1 DEGs for each cluster in scRNA-seq dataset and cell type annotation.

Table S2 Differentially expressed genes in TDP-43 K181E mutant organoids.

Table S3 Gene Ontology analysis of down-regulated TDP-43 binding RNAs identified by eCLIPseq.

Table S4 RNA binding sites detected by eCLIPseq.

Table S5 Detected alternative splicing events in homozygous K181E organoids.

Table S6 Top 20 predicted off target sequences for the sgRNA used in CRISPR editing

Table S7 DEGs for PRDM2 KD in motor neuron and Gene Ontology analysis.
